# Optimizing Predictive Metrics for Human Reading Behavior

**DOI:** 10.1101/2023.09.03.556078

**Authors:** Kun Sun

## Abstract

Expectation and memory have been found to play crucial roles in human language comprehension. Currently, the effects of both expectation and memory can be estimated using computational methods. Computational metrics of surprisal and semantic relevance, which represent expectation and memory respectively, have been developed to accurately predict and explain language comprehension and processing. However, their efficacy is hindered by their inadequate integration of contextual information. Drawing inspiration from the attention mechanism in transformers and human forgetting mechanism, this study introduces an attention-aware method that thoroughly incorporates contextual information, updating surprisal and semantic relevance into attention-aware metrics respectively. Furthermore, by employing the quantum superposition principle, the study proposes an enhanced approach for integrating and encoding diverse information sources based on the two attention-aware metrics. Metrics that are both attention-aware and enhanced can integrate information from expectation and memory, showing superior effectiveness compared to existing metrics. This leads to more accurate predictions of eye movements during the reading of naturalistic discourse in 13 languages. The proposed approaches are fairly capable of facilitating simulation and evaluation of existing reading models and language processing theories. The metrics computed by the proposed approaches are highly interpretable and exhibit cross-language generalizations in predicting language comprehension. The innovative computational methods proposed in this study hold the great potential to enhance our understanding of human working memory mechanisms, human reading behavior and cognitive modeling in language processing. Moreover, they have the capacity to revolutionize ongoing research in computational cognition for language processing, offering valuable insights for computational neuroscience, quantum cognition and optimizing the design of AI systems.

## 1 Introduction

Cognitive and behavior sciences have embraced computational methodologies to unravel the intricacies of human cognition and behavior, fueled by the success of large-scale deep learning models and the availability of vast datasets in cognitive and neural processing domains (Guest and Martin, 2021; Armeni et al., 2017; Crocker, 2012; Lee et al., 2019). The endeavor to comprehend how humans process language encompasses the utilization of meticulously crafted computational models, which simulate intricate cognitive processes and delineate a sequence of processing stages. These stages are systematically correlated with observed behaviors and neural responses, ultimately yielding quantitative insights into the underlying mechanisms governing language comprehension and usage.

Two theories concerning reading and language comprehension have been recently advanced in cognitive sciences, yielding notable influence. The first theory is **expectation**-based. Considering the pivotal role of expectations in language comprehension and processing (Clark, 2013; Huettig, 2015; Kuperberg and Jaeger, 2016), various computational models have been proposed to simulate the word predictions in a given context, with metrics like word **surprisal** (Hale, 2001; Levy, 2008) quantifying transitional probabilities between words. Expectation-based theories postulate that processing difficulty is largely driven by how predictable the upcoming linguistic material is in its context, and surprisal has been shown to be a strong predictor of the behavioural and neural processing measures of processing difficulty. The second theory is **memory**-based. In contrast, memory-based models (Baddeley, 2010; Lewis et al., 2006) posit that processing challenges result from constrained storage of prior context, as well as to retrieve and incorporate them with new input. Under this hypothesis, the semantic relatedness between a target word and its contextual words is crucial for word prediction. Memory-based approaches allow for interpretation of words based on established computational semantic relatedness, encompassing concepts like semantic vectors and **semantic similarity** (Mitchell and Lapata, 2010; Hollis and Westbury, 2016). The other computational memory-based models, such as dependency distance (Gibson, 1998; Liu, 2008), can also explain language comprehension difficulty.

Numerous studies have demonstrated the significant roles of expectation and memory models in language comprehension and processing, each contributing to individual effectiveness. However, there is still much room for improvement and refinement for their methods and their comprehensive integration has been seldom explored. The following provides a brief overview of their attempts and potential problems respectively.

Word predictability, similar to word probability (Staub, 2015), plays a crucial role in language comprehension and processing. It is rated by humans and is closely related to word surprisal, a computable metric that can be applied to large text datasets. Researchers have developed diverse methodologies for the automated computation of word probability and word surprisal, employing n-gram language models or neural language models. Much research has compared various language models that could help compute word surprisal more precisely (Wilcox et al., 2020; Hale et al., 2022). Nonetheless, language models merely serve as a facilitating tool in the computation of surprisal. Moreover, several methods have been proposed to enhance surprisal effectiveness. For instance, some research incorporated context information into surprisal to generate weighted surprisal (Sayeed et al., 2015; Russo et al., 2020). However, such methods are pretty complex and unexplainable, and the computational cost is also expensive. Without statistical comparison, there is little evidence to show the weighted surprisal outperforms surprisal. Evidently, researchers realized that a pivotal avenue for enhancing computational models in cognitive science lies in the effective incorporation of contextual information. However, the effectiveness of these endeavors has been seriously constrained by a range of factors. To effectively incorporate contextual information, we have to adopt a new perspective, taking further inspiration from specific empirical findings. For example, many studies found the effect of the predictability of the next word (i.e., “n+1” word) on processing the target word (Kliegl et al., 2004; Kennedy et al., 2013). It reveals that the contextual information on surrounding words’ surprisals actually impact the target word. This further suggests the potential utility of incorporating the surprisal information from contextual words. However, a challenge remains in optimizing a quantitative framework to integrate surprisal information from contextual words. The present study presents an innovative “attention-aware” approach designed to address this concern. Further details are elaborated in the following description.

On the other hand, a wealth of empirical research has discovered the effect of semantic plausibility in reading and language comprehension (Rayner et al., 2004; Hohenstein et al., 2010; DeLong et al., 2014; Veldre and Andrews, 2016). These findings emphasize the significant influence of the semantic connection between a target word and its neighboring words on the processing of that target word. Similar to word predictability, many of these studies have depended on human rating techniques to estimate semantic relatedness, which is then used to evaluate semantic plausibility. However, relying on human ratings becomes unfeasible when handling extensive text volumes. Thankfully, recent advancements in natural language processing (NLP) and artificial intelligence (AI) provide efficient methods for computing contextual semantic relevance (Sun et al., 2023a). Moreover, contextual semantic information can be linked with short-term memory, a notion supported by empirical investigations into memory systems like the visuospatial sketchpad and phonological loop (Baddeley, 2000; Postle, 2006; Baddeley, 2010). More holistic methods for computing contextual semantic relevance have been proposed (Sun et al., 2023a; Sun et al., 2023b; Sun, 2023). However, in terms of human forgetting mechanism (Murre and Dros, 2015), the current study proposes a method to compute contextual semantic relevance by considering the weights which are determined by the position distance between the target word and its surrounding words (Fig. 3A (a)(b)). Meanwhile, the following word information can be incorporated. Such strategies allow a more effective and reasonable incorporation of the contextual information. The approach used for this purpose is referred to as “attention-aware” (Sun et al., 2023b) (Fig. 2A, Table 1), inspired by attention architecture in Transformer (Vaswani et al., 2017) and the human forgetting mechanism (Murre and Dros, 2015). Meanwhile, the attention-aware approach can be applied to enhance the optimization of word surprisal by incorporating the information of surrounding words’ surprisals.

**Figure 1:**
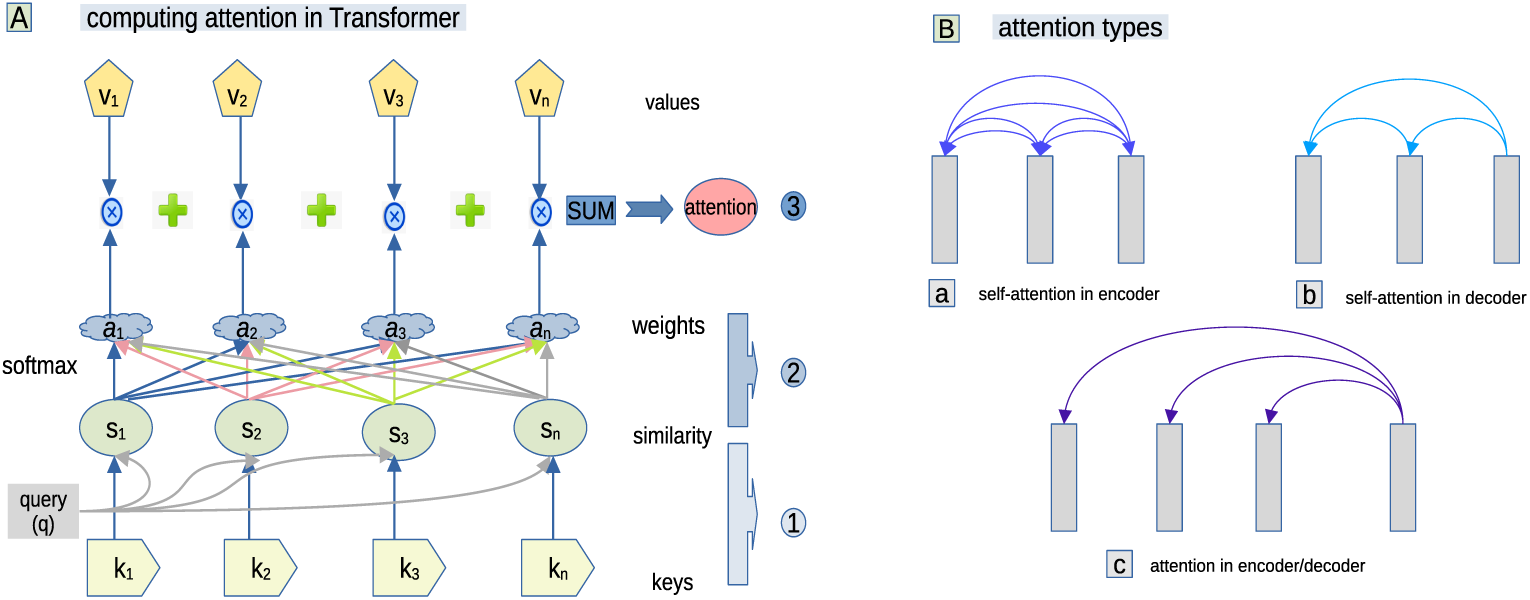
Attention mechanism in Transformers

**Figure 2:**
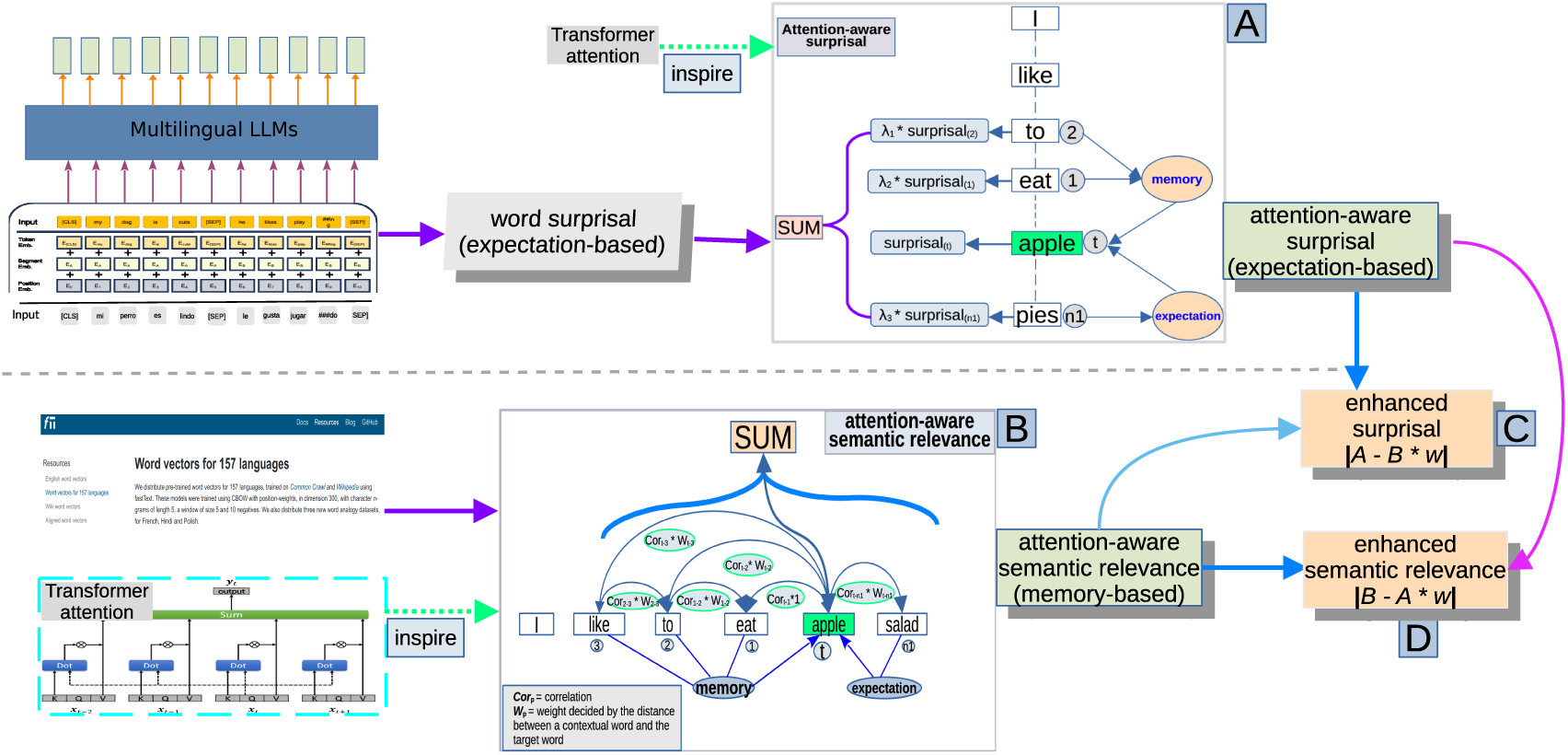
The methods of computing attention-aware measures and enhanced measures. (Panel A illustrates the computation of attention-aware surprisal, while Panel B demonstrates the computation of attention-aware semantic relevance metrics. Panels C and D depict the use of weighted superposition to compute two distinct types of enhanced metrics, each in its own respective context. Here we used three Multilingual LLMs (i.e., m-BERT, XLM, and mGPT) to compute word surprisals.)

**Table 1:**
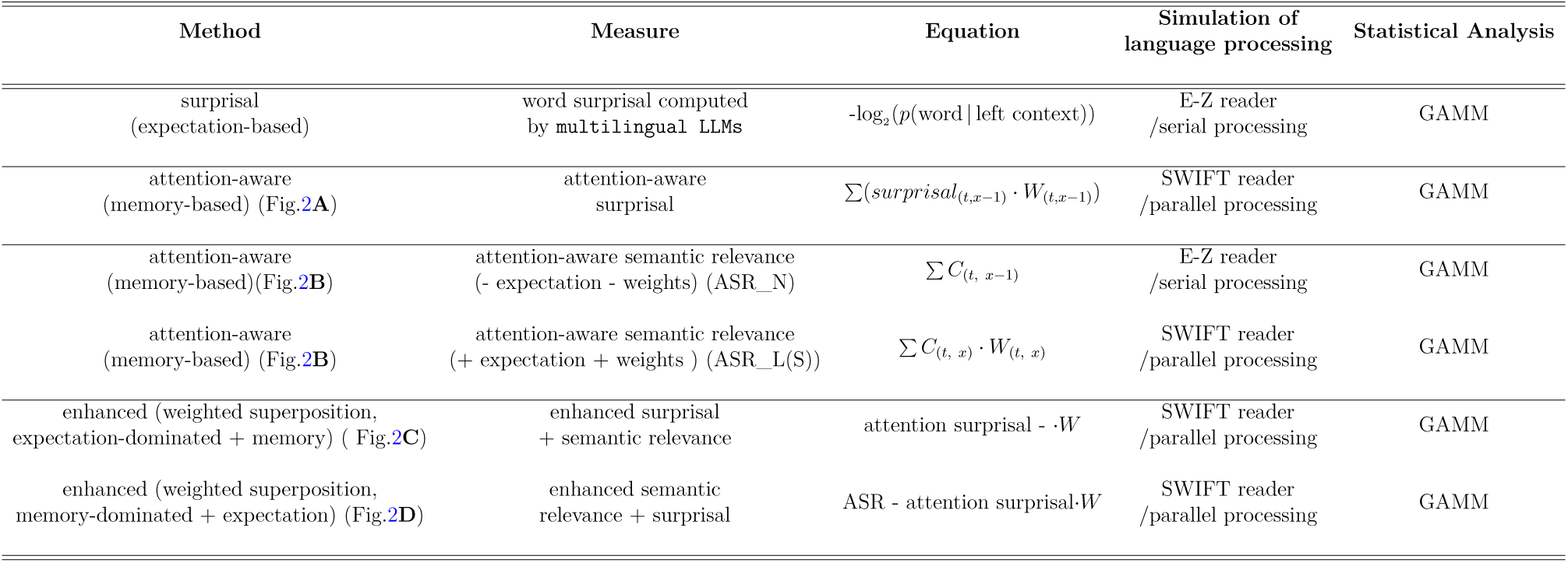
The methods and metrics used in this study

The comprehensive integration of expectation and memory effects has been seldom explored. Prior attempts to combine expectation-based and memory-based effects, like surprisal and dependency length (Futrell et al., 2020; Hahn et al., 2022), have lacked a feasible computational implementation. The pursuit of a practical solution for effectively merging these models has remained elusive. To tackle this issue, the present study proposes another novel approach, termed the “enhanced” approach. This approach skillfully integrates surprisal with semantic relevance, resulting in the creation of more robust and reliable metrics (“enhanced” metrics). The enhanced approach combines the two metrics using a weighted superposition technique, a method commonly employed in the fields of physics and quantum sciences (Gudder, 1970; Rudin, 2017; Koopmann et al., 1989; Busemeyer and Wang, 2015). The superposition of the two metrics may be equivalently construed as a superposition of quantum state. These metrics are designed to better explain and predict language comprehension and processing.

Besides expectation and memory theories, various theories have been put forth to elucidate human reading mechanisms (Kliegl et al., 2006; Clifton Jr et al., 2016). Eye-movement control models explain the sequence of fixations and saccades that transpire during reading. The E-Z Reader model, for instance, suggests that lexical processing unfolds in a sequential manner, with parafoveal effects being a rare occurrence (Reichle et al., 2003). On the other hand, the SWIFT model posits that such effects are commonplace, attributing this to the parallel processing of words (Engbert et al., 2005). The E-Z Reader model lends support to the concept of serial processing, whereas the SWIFT model champions the idea of parallel processing (Reilly and Radach, 2006; Snell and Grainger, 2019). These hotly debates have persisted for almost two decades without reaching a definitive resolution. These discussions have been driven by numerous empirical experiments conducted with small-sized datasets. This is because these theories/models in question (actually many cognitive models/theories) are constrained by computational limitations and have restrictions in generating metrics for evaluating larger datasets. However, the approaches we proposed, along with the metrics generated, have the capability to simulate these theories/models using extensive datasets. Further, we are interested in determining whether these metrics can quantitatively evaluate the relative performance of each theory or model using large-scale datasets.

To achieve these goals, we plan to implement the newly proposed approaches to generate a range of metrics, evaluating their predictive efficacy across multiple languages beyond English. Most language cognition research centers on English, but its applicability to other languages remains uncertain. This bias can lead to over-generalizations (Blasi et al., 2022). We will carry out multi-lingual investigations to test cross-language generalizations of these metrics. Such cross-lingual investigations would need to draw on multi-lingual databases of language cognition and use effective multi-lingual computational tools. Fortunately, such databases and computational tools are available. For instance, multi-lingual large language models (LLMs, e.g., m-BERT, Devlin et al. (2018); XLM, (Conneau et al., 2019); mGPT, (Shliazhko et al., 2022)) are capable of processing and understanding text in multiple languages. Multilingual LLMs can estimate word surprisal precisely respectively for multiple languages. There are also multilingual databases on human language processing. For instance, the Multilingual Eye-movement Corpus (MECO) is a collection of eye-tracking data that has been collected from participants reading texts in 13 languages (Siegelman et al., 2022). The MECO enables us to conduct experiments in which these metrics are tested to show their predictability for the eye-movements in multiple languages.

The present study promises to enhance our comprehension of human reading behavior and language processing intricacies. The approaches and metrics proposed can revolutionize ongoing research in computational cognition for language processing, providing insights for computational neuro-science and quantum cognition, and enhancing AI system design.

## 2 Methods

### 2.1 Testing datasets

There are extensive multilingual databases available for cognitive and neural language processing, and utilizing these databases to test model predictions is both cost-effective and yields reliable results. Eye-tracking data is particularly useful in assessing computational language models, as fixation duration is closely linked to the cognitive effort required for processing. Although English dominates eye-tracking research, the Multilingual Eye-tracking Corpus (MECO) (Siegelman et al., 2022), which comprises 13 languages, provides a more diverse range of languages. In the MECO corpus, participants in each language were presented with 12 texts, each containing 10-15 sentences, resulting in 2000 tokens per language (2000 x 13 = 36000 tokens). The texts were encyclopedia entries on various topics with similar complexity and readability across the languages. Each language yielded approximately 70,000 to 80,000 recorded tokens, providing a sufficient amount of eye-tracking data. The MECO corpus was chosen for several reasons, including its inclusion of databases in 13 languages with similar text stimuli, large numbers of native speakers for each language, and a sufficient amount of eye-tracking data. The present study focuses on three specific eye-movement measures as response variables: first duration, total duration, and gaze duration. First duration refers to the duration of only the first fixation on the target word, total duration refers to the summed duration of all fixations on the target word, and gaze duration refers to the sum of the fixation durations before the target word is exited to the right or left during first-pass reading.

Using the MECO as the testing data involves additional considerations. The simplicity and directness of eye-movement data (i.e., the data on fixation duration) lends itself well to conducting advanced statistical analyses for comparing the efficacy of different metrics in predicting eye-movements. In contrast, EEG or fMRI data, for example, are notably more intricate and subject to diverse manipulation methods, making them less amenable to straightforward statistical analysis when evaluating the predictive performance of these computational metrics on such data. Moreover, previous cognitive studies may have been distorted by a research focus on the English language, leading to unsupported over-generalizations based on features and mechanisms specific to English. Conducting wider studies can help avoid such distortions and ensure that our metrics are based on properly founded scientific **generalizations**.

### 2.2 Training datasets

Three multilingual transformer-based large language models (LLMs) were employed to compute word-level surprisal for various languages. Multi-lingual BERT Devlin et al. (2018) is one influential multilingual language model. We use the state-of-the-art BERT model (i.e., multilingual-bert-uncased, abbreviated as m-BERT) because it can be consistently applied in different languages in order to compute surprisal for word in a text of that language.

A BERT model was trained pretrained with a masked language modeling (MLM) objective, and therefore it can be used to compute word probabilities, from which we can estimate word surprisal scores. In order to make verification, another influential multilingual LLM (i.e., XLM (xlm-roberta-large) (Conneau et al., 2019)) was utilized to compute word-level surprisal for multiple languages. Similarly, XLM was also pretrained with the Masked language modeling (MLM) based on more than 100 languages.

The third multilingual LLM is mGPT (Shliazhko et al., 2022). mGPT is a multilingual variant of the GPT model, trained on 61 languages. It is an autoregressive model with 1.3 billion parameters, trained using Wikipedia and the Colossal Clean Crawled Corpus. The model reproduces the GPT-3 architecture using GPT-2 sources and a sparse attention mechanism. The mGPT model, a multilingual adaptation of the GPT architecture, is capable of calculating word probabilities and word surprisals across multiple languages. Unlike the MLM approach used in BERT, which predicts masked words based on their context, GPT employs an autoregressive language modeling technique. This method involves predicting the probability of the next word in a sequence, given all the previous words.

Note that the mGPT model from STOA does not support the Norwegian language. Consequently, we need to establish two distinct groups for comparing the predictability of metrics computed by these LLMs. The first group employed m-BERT and XLM to process the entire testing dataset of MECO, including Norwegian. This provided us with a comprehensive understanding of their performance across all languages in the dataset. The second group involved a comparison between m-BERT, XLM, and mGPT. However, in this case, the testing dataset (i.e., MECO) excluded Norwegian, given mGPT’s lack of support for this language. This allows us to evaluate the performance of these models on a level playing field. The results corresponding to these two different scenarios are reported in separate sections, ensuring a clear and concise presentation of our findings.

However, unlike computing surprisal, we used multilingual fastText pre-trained databases (Mikolov et al., 2017; Grave et al., 2018) in each language to help compute word-level attention-aware measures of contextual semantic relevance. These pretrained databases of word embeddings for multiple languages are available at https://fasttext.cc/docs/en/crawl-vectors. html. Using contextualized embeddings generated by LLMs is a viable option. However, employing these contextualized embeddings for computing attention-aware metrics can lead to an issue: it becomes unclear whether the predictability of attention-aware metrics is a result of the attention-aware approach itself or the integration of contextual embeddings to capture contextual information. To mitigate this potential ambiguity, opting for static pre-trained word embeddings emerges as a prudent decision. Moreover, the word frequency information across languages in the present study is obtained from https://opus.nlpl.eu/index.php (Lison et al., 2018).

### 2.3 Methods of computing surprisal

**BERT-based surprisal** can be formalized as the following equation.

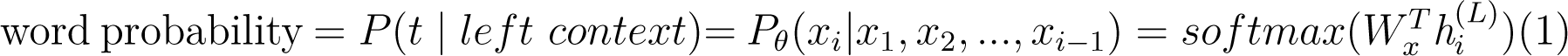

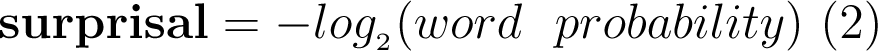

“*h*_*i*_^(*L*)^ ” is the vector for the *i*th position word. “x_1_, x_2_, …, x*_i−_*_1_” are the other words on the left context of the target word (i.e., “x*_i_*”). “t” here also refers to target word. “W*_x_*” is the weight matrix for all words in this window. *θ* is the model parameter.

The following details how LLMs were implemented to estimate word surprisal. We first loaded m-BERT (or xlm) and tokenizer from the Hugging Face Transformers library. We then defined an input text sequence with a [*MASK*] token (i.e., MLM), which the model will try to predict. Next, we used the trained model to generate a probability distribution over the possible next words for the masked token in the input sequence. We did this by passing the input tensor to the model, and then extracting the logits and softmax probabilities for the masked token. We then found the word with the highest probability in the distribution to use as the next word for the surprisal calculation. We computed the surprisal value as the negative log probability of the actual next word in the distribution.

Contrary to the MLM approach, mGPT employs an autoregressive language modeling technique. This method focuses on predicting the probability of the next word in a sequence, given all the preceding words. Once these word probabilities are computed, we can readily calculate the word surprisal. Additionally, calculation of word surprisal takes place within a sentence. Notably, the first word in a sentence holds no surprisal value as it lacks a preceding context.

### 2.4 Computing attention-aware metrics

Before delving into the computation of attention-aware metrics, it is essential to provide a step-by-step explanation of how to compute attention in transformers, which are the state-of-the-art deep learning model (Vaswani et al., 2017). This allows us to highlight the similarities and differences between attention in Transformers and attention-aware adopted in the present study and better understand their underlying mechanisms.

We compute transformer attention scores in four steps, which is shown in Fig. 1A. Step 1 is to create input vectors X, and combine the input vectors with the positional encoding. Step 2 is to derive three vectors: query, key and value by multiplying 3 the initialized weight matricies W with X. q, k, v are three abstractions, or linear transformations from the same source. They will be useful in computing attention. Those initialized weights will be trained and optimized. In step 3, we can select a single token from the Query vector, such as q, and compute the relevance or alignment between this query and all other keys, including itself, using a dot product. Step 4 allows us to use softmax to normalize the relevance or alignment scores, resulting in attention weights. These values, ranging from 0 to 1, indicate where to focus when processing the token. In step 4, we compute a weighted sum of the values using these attention weights to get the output/ context vector for the token q. Steps from 3 to 5 can be represented by a mathematical equation, shown in Equation (3).

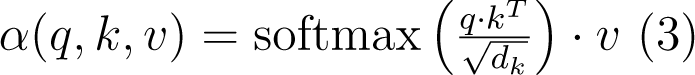

Here, “q” represents the query, “k” represents the key, and “v” represents the value. These are matrices derived from the input embeddings. The soft-max function normalizes the weights, and d*_k_* is the dimension of the key vector, which is used for scaling. The process repeats for every token in the input sequence. Finally, we obtain a context vector from the entire input vector X. The context vector has the same shape as input vector X but contains rich contextual information. Those context vector will be fed to the next layer.

There are various types of attention in transformers, as shown in Fig. 1B. The attention layer measures the similarity between the query and the key using a score function *α* which returns scores *a*_1_ , . . . , *a_n_* for keys *k*_1_ , . . . , *k_n_* given by. The score function, *α*(*q, k*), can take various forms, which has given rise to a number of attention mechanisms, show in Equation (3) (Vaswani et al., 2017). Once the attention weights (*α*) are computed, we multiply them by the value vector *v_i_* to obtain an updated representation for embedding.

The main idea behind attention in Transformers is to associate each element in the input sequence, such as each word in a sentence, with a weight that determines its importance in computing the context vector. This allows the attention mechanism to link the target word with all other words in the input sequence. However, the embeddings incorporating attention information in Transformer are trained and optimized in neural networks, making them become **uninterpretable**, so we **cannot** use the embeddings in transformers directly.

While the attention architecture in transformers is inspiring, it is necessary to **customize** it to meet specific requirements for cognitive studies (i.e. language comprehension and processing). For instance, we need to significantly reduce the window size to between 5 and 6 words and use the position distance between the target word and its contextual words as weights instead of average weights and positional embeddings. By combining the contextual semantic relevance method (Sun et al., 2023a) with these modified attention mechanisms, we present an improved and more reliable “attention-aware” method. To illustrate the attention-aware approach, let us consider attention-aware semantic relevance. In this case, we sum up all the values of semantic similarity (cosine or correlation) between the target word and each of its surrounding words (5-6 words) and the correlation among these surrounding words. Meanwhile, we consider the distance between the surrounding words and a target word as weights which are used to multiply with the corresponding semantic similarity values and adding the expectation effect of the next word. Further, the association between the target word and its preceding words, and the relevance among the preceding words incorporate the memory-based information. After the following word as the expectation effect is taken into account, the method incorporate both memory-based information and expectation-based effect. In this sense, this method can incorporate semantic relatedness as derived from the context more comprehensively, that is, the method can incorporate the contextual information more fully. Our method plays a similar role in that of the attention mechanism in Transformer capturing contextual information. We therefore define it “attention-aware” approach. Equation (5) represents the semantic relevance computed by the attention-aware approach in the current study.

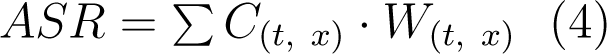

Here “ASR” is the abbreviation of “ attention-aware semantic relevance”, and “t” represents target word, “x” for contextual words (the preceding three words and the following word). “*C* ” represents correlation, and “*W* ” for weights.

Fig. 2B shows the computation of the attention-aware semantic relevance (ASR) for the target word “apple”, which is surrounded by “like”, “to”, “eat”, “pies”. To compute attention-aware semantic relevance, we assign each word pre-trained word embedding. The words preceding the target word (e.g., ‘like’, ‘to’, ‘eat’) form a short-term memory stack, simulating how readers store memory of previously encountered words and their meanings. Using word embeddings, we employed the *correlation* to obtain a value to represent the semantic relevance between two words in this stack, and the value indicates the memory retained and the semantic relatedness within the context. We also included the next word (e.g., ‘pie’) to represent the expectation information and expand the contextual window. The semantic relevance between the target word and this next word is calculated to represent such information. Weights are assigned based on the distance between the target and contextual words. Specifically, we multiply each correlation value by a weight factor that ranges from 0 to 1, with closer words receiving higher weights. Words that are closer to the target word are assigned higher weights. For example, “Cor*_t_* _3_” (correlation between “apple” and “like”, here “t” and ”1” representing target word and the left first close word) is multiplied by 2/3, and “Cor*_t_* _2_” (correlation between “apple” and “eat”) is multiplied by 3/4. The correlation value for “like” or “to” is reduced because they are not directly adjacent to the target word “apple”. “Cor_2_*_−_*_3_” (correlation between “like” and “to”) is multiplied by 2/3, and “Cor_1−2_” (correlation between “to” and “eat”) is multiplied by 2/3. The correlation value for “apple” and the following word “pies” is also reduced by a factor of 1/3 since we believe that the expectation effect is not as strong as memory-based information. After each correlation value is multiplied by its corresponding weight, we summed up all the values to obtain the “attention-aware semantic relevance ” value. The current study adopts two sets of weights, large and small ones (i.e., ASR_L(with small weights), and ASR_S (with large weights)).

Equation (5) represents the case that the following word and weights are not considered, and it is

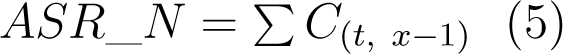

Here “ASR_N” is the weak version of attention-aware semantic relevance (“N” here refers to no weights). “t” represents target word, “x-1” for contextual words (the preceding three words but excluding the next word). “C” represents correlation.

“The attention-aware semantic relevance (ASR)” represents the degree of semantic relatedness of the target word with its context. Shown in Fig. 2B, ASR for the target word ‘apple’ indicates the semantic relationship between ‘apple’ and its surrounding context, reflecting the degree of their semantic connection. When the value of ASR is large, it means that the target word is closely related with the context or highly plausible with the context. With a larger ASR, there is an implication of enhanced capacity to retain contextual representations. This, in turn, facilitates the retrieval and integration of these representations with the new input, such as the target word “apple”, in accordance with the memory-based theories. Consequently, the processing of the target word becomes less challenging, leading to potentially shorter reading durations. On the other hand, when the value of ASR is small, it suggests a diminished semantic connection between the target word and the context, or a reduced contextual plausibility. In terms of memory-based theories, this lower ASR value also signifies a weaker ability to store contextual information, thereby making retrieval and integration with the target word less straightforward. As a result, processing difficulty intensifies, necessitating more time.

Adopting the similar attention-aware approach, we can create “attention-aware surprisal”. Looking at the preceding two words and the following word, we add up the surprisal values of the two preceding words and the one of the following word. However, weights will be multiplied with each of the three surprisal values. Otherwise, the direct summation with the surprisal of the target word does not work. It means that the contextual surprisal will influence the surprsial of the target word. However, their effects are fully and immediately taken on the target word. Similarly, the weights will changed with the position distance between the surrounding words and the target word. Its equation is shown as in (6).

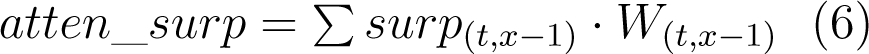

Here “atten_surp” refers to “attention-aware surprisal”. “surp” is for “surprisal”. “t” represents target word, “x-1” for contextual words (the preceding three words but excluding the next word). “*W* ” represents weights.

We now detail how to calculate “attention-aware surprisal”, as shown in Fig. 2A. First, we calculated the surprisal values of the preceding two words (using m-BERT), “to” and “eat”, as well as the following word, “salad”. Then, we multiply the surprisal values of the two words in the neighborhood by 1/3 (weight) each, while the surprisal value of the far preceding word “to” is multiplied by 1/4 (weight) because of its greater distance from the target word. Unlike semantic relevance, we only include the preceding two words in computing attention-aware surprsial. The term “attention-aware surprisal” refers to contextual surprisal, which entails incorporating the surprisal of the target word with the associated surprisal values of contextual words. The information on contextual surprisal influences the processing of the target word by facilitating memory retrieval and integration. In the case of the same word, a higher attention-aware surprisal signifies larger surprisal values for the contextual words. This amplified impact heightens the processing difficulty. Conversely, a lower attention-aware surprisal for the same word indicates smaller surprisal values for the contextual words, resulting in a diminished impact and reduced processing difficulty. Note that all attention-aware metrics are computed within a sentence.

The following explains why weights and next word information are taken in the attention-aware approach. The attention-aware approach employed in this study is fundamentally memory-based and has potential implications related to human memory processes. The preceding words in a window, as illustrated in Fig. 2, simulate a memory stack because readers retain their memory of previously encountered words and their meanings (4-5 words in a stack). The weights assigned to compute attention-aware metrics are dependent on the distance between the target word and its surrounding words. Closer words receive higher weights, while more distant words receive lower weights. In this study, we utilized a window size of 4-5 words, and it could reflect the shape of a forgetting curve, that is, a representation of memory retention decline over time, where retained information halves after each day within a span of several days (Loftus, 1985). In this window stack, words closer to the target word resemble the initial days in the forgetting curve, while more distant words resemble the latter days. To simulate human forgetting mechanism, we allocated larger weights to the closer words and smaller weights to the distant wordsand the similarity between human forgetting mechanism and attentional weights adopted in the current study is illustrated in Fig. 3A. The use of different weights is effective in integrating information on the varying contributions of contextual words, and the various weights can also encode information on word order.

By considering both preceding and the following word (“n+1”) as potential sources of contextual information, the attention-aware approach (i.e., ASR, attention-aware surprisal) enables a wider window size it can simulates the SWIFT model (or parallel processing, Engbert et al. (2005); Snell and Grainger (2019)). The attention-aware metric (i.e., “ASR_N) without considering the “n+1” word can stimulate E-Z reader or serial processing (Reichle et al., 2003). This is illustrated Fig 3B.

**Figure 3:**
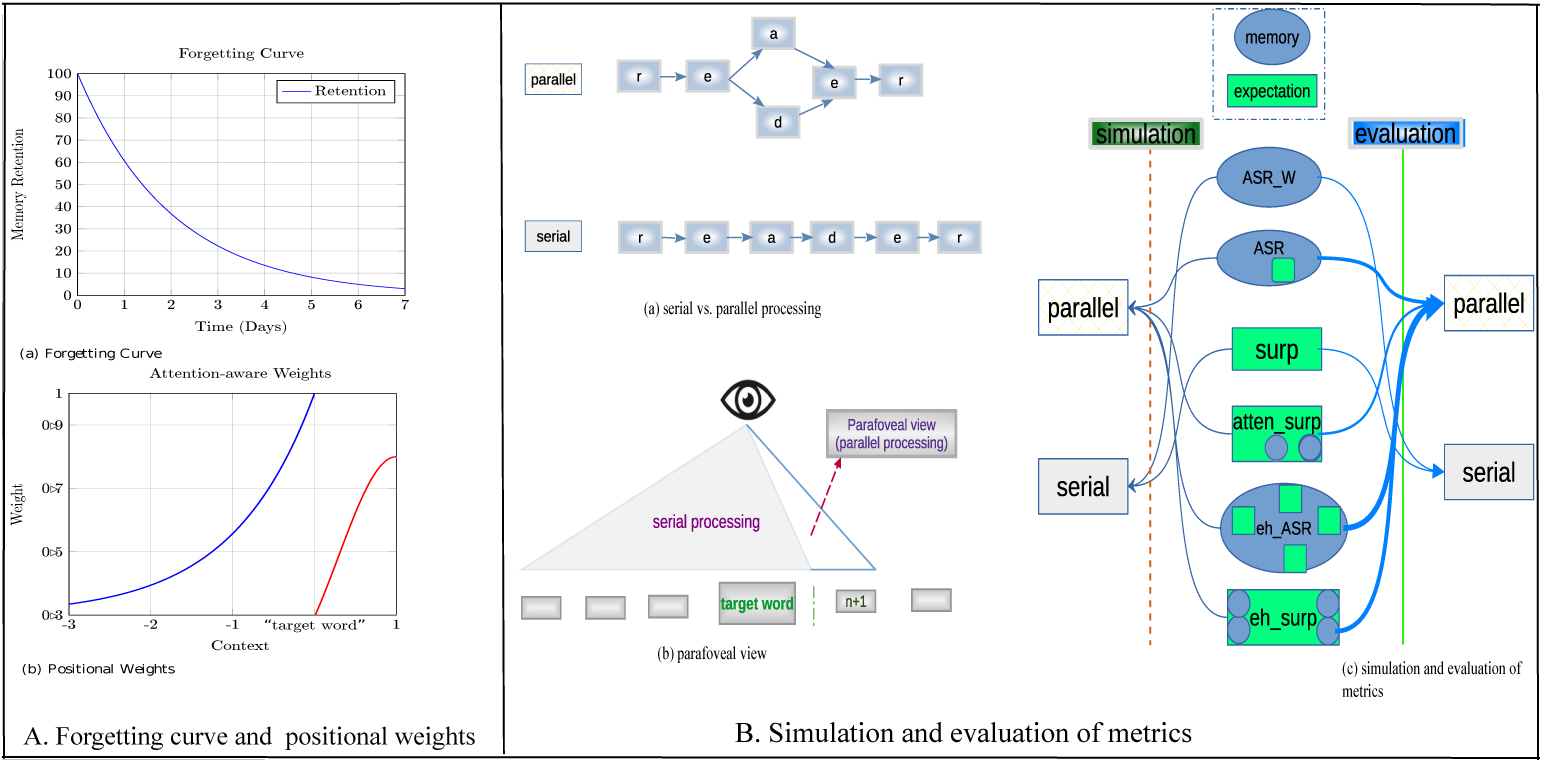
Memory mechanisms and quantitative simulation/evaluation. (Panel A shows similarities between human forgetting curve (a) and the attention-aware weights(b) where both decay as time goes. Panel B illustrates serial vs. parallel processing in language comprehension (a), and preview in reading (b), as well as how our metrics stimulate these language processing theories quantitatively and evaluate them for largescale datasets (c).)

In brief, the attention-aware approach is both computational and fundamentally memory-based, facilitating effective incorporation of contextual information and enabling it to achieve memory storage, retrieval, and integration, shown in Fig. 6. The attention-aware approach can allow metrics to capture long-term dependencies and store short-term memory, much like the transformer architecture. The attention-aware method is highly interpretable and versatile, and can be applied to optimize various metrics, enabling them to capture contextual information.

### 2.5 Computing enhanced metrics

The attention-aware metrics have significant implications for explaining and predicting language processing. However, by effectively incorporating the expectation and memory effects, we created an “enhanced approach”. The enhanced approach is grounded in the superposition principle, a concept extensively applied in quantum physics, engineering, and cognitive studies (Gudder, 1970; Rudin, 2017; Koopmann et al., 1989; Busemeyer and Wang, 2015).

The superposition principle is derived from the algebraic summation of all stimuli at a specific point in space-time. The mathematical framework for interference is constructed on the linear superposition of waves, a principle vividly illustrated in the properties of the Fourier transform. Linear combination is a remarkably intriguing concept that seems to apply widely. The concept of linear combinations is central to linear algebra and related fields of mathematics. When a system appears to be undergoing superposition, what actually occurs is the cumulative convergence of the weighted causative factors behind two or more similar phenomena, culminating in an outcome that is meticulously scaled in proportion. The two weighted superposition equations can be summed up as a linear combination function:

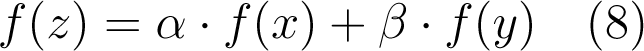

Here *α* and *β* are weights.

Following the weighted superposition principle, we can generate two types of “enhanced” metrics. First, “enhanced surprisal” (enhanced surp) merges the attention-aware surprisal and attention-aware semantic relevance. On the other hand, the “enhanced semantic relevance measure” (enhanced ASR) is obtained by incorporating the attention-aware surprisal information into attention-aware semantic relevance. Fig. 2C, D show the computation of enhanced surprisal and enhanced semantic relevance, respectively. The enhanced method diverges slightly from the traditional superposition principle in that it employs weights to reduce the value of one metric before combining them. Without this adjustment, the direct summation of the two metrics would result in destructive interference. By incorporating a weight, constructive interference is achieved, as depicted in Fig. 4A(a). Conversely, when attention-aware surprisal and attention-aware semantic relevance are summed without modification, it implies that readers consider both equally, akin to waves 180° out of phase. This leads to a destructive interference effect, as illustrated in Fig. 4A(b).

**Figure 4:**
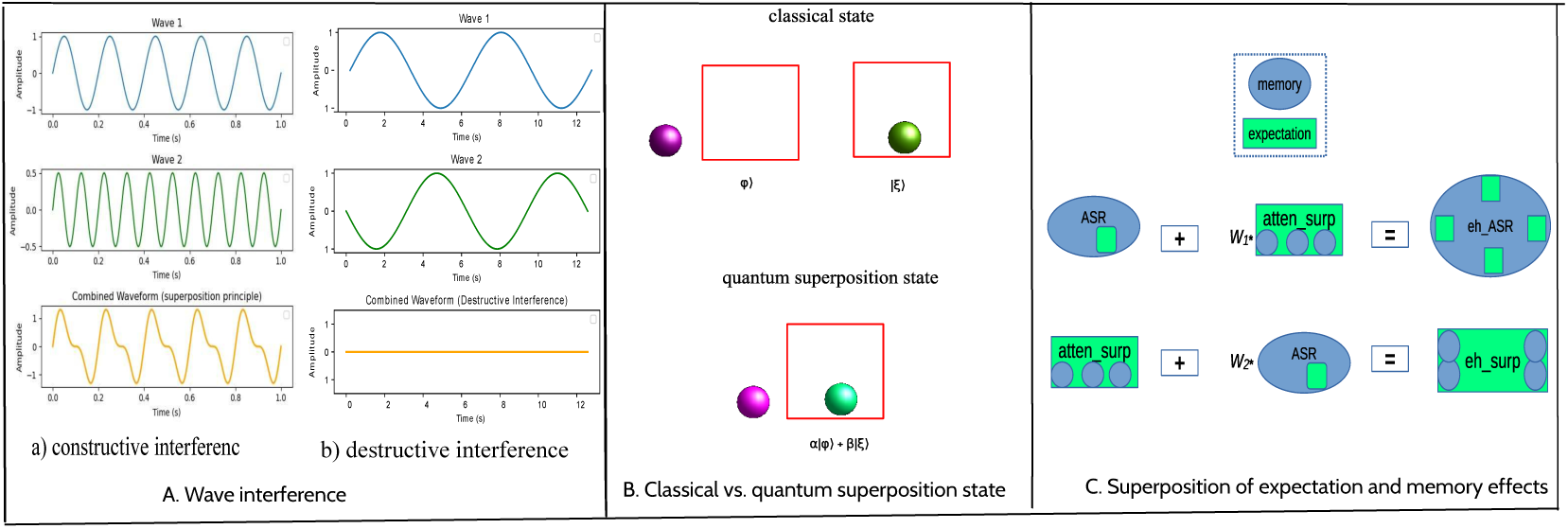
Superposition principle and enhanced metrics. (Panel A depicts two instances of wave superposition: a) leading to constructive interference, and b) resulting in destructive interference. Panel B illustrates the quantum superposition state, which is different classical state. The quantum system is only in one state at a time, and this state can be expressed as a superposition of other states. Panel C demonstrates the utilization of weighted superposition for calculating two distinct types of enhanced metrics. The blue oval shape signifies memory, while the green square shape represents expectation. This differentiation is why the term ‘ASR’, along with subsequent word information, incorporates a smaller expectation symbol, whereas ‘atten_surp’ includes two memory symbols (also see Fig. 3B(c)). Enhanced metrics encompass both expectation and memory effects through weighted superposition. However, each enhanced metric has its distinct dominant effect (but inserted with subdominant factors). After the superposition, the size of the enhanced metric becomes larger, signifying that their abilities also become stronger.)

The “enhanced” approach, which encompasses both memory and expectation effects while encoding word order information, can simulate parallel processing, shown in Fig. 4C. In Figure 4C, the blue oval shape signifies memory, while the green square shape represents expectation. This distinction is why ‘ASR’, includes a smaller expectation symbol, whereas ’atten_surp’ incorporates two memory symbols. “Enhanced metrics” include both expectation and memory effects. This is why each enhanced metric symbol encapsulates these effects so prominently. “Enhanced surprisal” is a form of surprisal that integrates information on semantic relevance, while “enhanced semantic relevance” is a measure of semantic relevance that incorporates information on surprisal. The following provides the equations for them respectively.

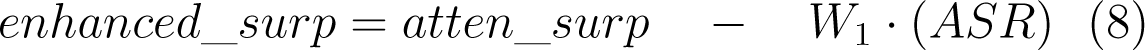

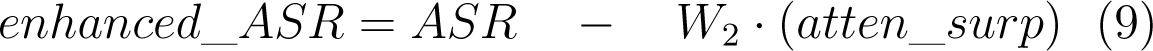

(Note: “*W* ” represents weight, and its value can be adjusted in terms of different corpora. The scope of two weights is as follows: 0.5 <*W*_1_ <0.7, optimal *W*_1_ = 2/3; 0.03 <*W*_2_ <0.1, optimal *W*_2_ = 0.02. For the current corpus, we found the optimal values. “enhanced_surp”= enhanced surprisal; “enhanced_ASR”= enhanced semantic relevance; “atten_surp” = attentionaware surprisal; “ASR”= attention-aware semantic relevance)

Weights are the key in the superposition method we adopted to compute enhanced metrics. However, two different weight values were used in the calculation. This is due to the fact that the two metrics have different levels of dependence on each other. Semantic relevance has a stronger effect, hence the weight of surprisal is relatively small in Equation (8). On the other hand, the weight of semantic relevance is much larger in Equation (9) because the effect of semantic relevance outweighs that of surprisal. We also provide a range of weights, which can be adjusted depending on the language and the task being analyzed. The different weights arise from the significant role that dominance plays in human behavior, cognition and neural activities (Demaree et al., 2005; Chen Zeng et al., 2022; Mascaro et al., 2023). One factor assumes dominance, while the others adopt a subdominant position, and their synergistic superposition yields exceptional outcomes. Although the two enhanced metrics are computed simply, they remain highly interpretable. Nevertheless, the range of appropriate weights has been obtained through our empirical experiments.

Our enhanced approach can be supported by quantum sciences. Being in a superposition state means that all potential measurement values maintain the capacity to appear at any moment. These potentials can engage with each other in a way reminiscent of wave interference, consequently modifying the final observed measurement value. Specifically, in the context of quantum cognition (Busemeyer and Wang, 2015), the reading state can be seen as a superposition of factors that have the potential to influence outcomes. These factors may interact, sometimes enhancing each other, leading to changes in observed measurements, like eye-movement data (e.g., fixation duration, fixation number, saccade length, skip etc.). We can also use quantum state to describe Equations (7), (8) and (9). A quantum state of a system is represented by a vector in a mathematical space known as a Hilbert space. The state vector is often denoted as |*ψ*⟩ and contains information about the system’s properties. The superposition principle states that if |*ϕ*⟩ and |*ξ*⟩ are two valid quantum states, then any linear combination of these states, *α*|*ϕ*⟩ + *β*|*ξ*⟩. The coefficients *α* and *β* determine the weight or probability amplitude of each state in the superposition. A Hilbert space is a mathematical space used to represent the quantum states of a system. In a Hilbert space, interference effects could occur when different quantum states are combined in a superposition and then interact with each other. Interference can lead to constructive or destructive interference, influencing the probability distribution of measurement outcomes. Finally, we can formulate the quantum superposition state using Equation (11) (Fig. 4B):

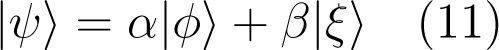

When a measurement is performed on a quantum system, the superposition collapses to one of the possible states. The eye-tracker could record the state as some given indicator. It also means that the enhanced metrics and reading processes could be interpreted from the perspective of quantum superposition.

In short, our enhanced approach is backed by linear combination, the weighted superposition principle, and quantum superposition. This empowers the incorporation of diverse sources for generating more resilient and robust metrics.

### 2.6 Statistical methods

To meet our goals of accurately predicting multilingual eye-tracking data, we utilized Generalized Additive Mixed Models (GAMMs) (Wood, 2017). GAMMs are effective in analyzing nonlinear effects and multiplicative interactions between variables, making them ideal for evaluating the predictability of semantic similarity. They are more flexible than traditional regression methods in modeling complex relationships between variables. Eye-tracking data is simpler to analyze statistically than EEG and fMRI data, which makes it an ideal choice for our study on naturalistic discourse reading. However, assessing model performance and comparing models can be challenging, and relying solely on correlations can be limiting. Fortunately, GAMMs are well-suited for comprehensive and precise assessments of model performance.

Comparing GAMM fittings, a larger model will have a higher likelihood due to more parameters. But, comparing loglikelihoods (or log-likelihood (LogLik)) directly is not suitable for differently sized models. Both AIC Akaike information criterion) and Bayesian Information Criterion (BIC) address this by penalizing model complexity. However, AIC is preferred for prediction quality, while BIC prioritizes parsimony, often favoring smaller models. Previous studies (Wilcox et al., 2020, Oh and Schuler, 2023) used ΔLogLik to analyze the predictability of surprisal estimated by various language models. However, LogLik may not be a good standard. Moreover, including random variables and significant predictors (control predictors) like “word length” and “word frequency” is crucial for accurate regression model evaluation. Failure to do so may result in suboptimal GAMMs or LMERs for studying reading time. Without optimization, there is a risk of encountering misleading issues that can undermine the validity of the results. In brief, the regression models in these studies (Wilcox et al., 2020; Oh and Schuler, 2023) may not be optimal because of lack of random variables or control predictor.

The present study included these control predictors (i.e., word length, word frequency) and random variable (i.e., “language”, “participants” explored as random smooth which includes random intercept and slope) in the GAMMs. Moreover, while certain studies have indicated that information regarding word length or word frequency in the preceding or following word can impact the processing of the target word (Kliegl et al., 2004), their effects appear to be comparatively weaker as control predictors when contrasted with the frequency and length of the target word (Schotter et al., 2012; Brysbaert et al., 2018). Hence, our GAMM fittings merely include the word frequency and word length of the target word as covariates to ensure robust control.

The performance of GAMM fittings was evaluated using difference in AIC (ΔAIC) between a base GAMM and a full GAMM served as a measure to assess the effectiveness of a computational measure. A smaller ΔAIC value indicates that the measure we are interested in estimates provide more accurate predictions of fixation duration compared to the baseline model. Table 1 provides an overview of the six metrics used in our analysis.

## 3 Results

### 3.1 Overall performance

This section reports how the surprisal, attention-aware metrics and enhanced metrics to predict the eye-movements on reading in the entire MECO (13 languages). Note that word surprisals were computed by m-BERT (Devlin et al., 2018) and XLM (Conneau et al., 2019) for 13 languages, respectively. mPGT Shliazhko et al. (2022) is not able to compute Norwegian which is one of 13 languages in the MECO, so we reported the results of surprisal estimated by mGPT in the next section.

Initially, we utilized Pearson’s correlation coefficient to investigate the relationships among these metrics. Our analysis revealed that surprisal and Attention Semantic Relevance (ASR) are fundamentally distinct metrics. Similarly, attentional surprisal and enhanced surprisal demonstrate significant differences from both ASR and enhanced ASR metrics. This pattern of distinctiveness is consistently observed across each analyzed language. Detailed findings are presented in the Appendix.

We then fitted 21 GAMM (generalized additive mixed model, Wood (2017)) fittings to analyze the seven metrics as predictors of three dependent variables (*total duration*, *first duration*, and *gaze duration*). The main predictor of our interest is modeled as a tensor product smooth. The GAMM fittings also include *word length* and *word frequency* as control predictors, modeled as tensor interaction, and *participant* as a random effect. An optimal GAMM fitting is formulated as: <u>duration∼te(word_length,log_wordfreq) +s(metric,language,bs=“fs”,m=1)+s(participant,bs=“re”),data=data</u> (s = tensor product smooth, te = tensor interaction, fs = random smooths adjust the trend of a numeric predictor in a nonlinear way, and it cover the function of random intercept and random slope, re = random effect is random slope adjusting the slope of the trend of a numeric predictor). In this GAMM equation, language and participant are treated as random variables. Our method leverages random slope and random intercept to fully assess the significance of the metrics of interest.

We used a threshold of *p*-value < 0.01 to determine the significance of variables in a GAMM fitting. The results of optimal GAMM fittings show that two types of surprisal predict first fixation, total fixation, and gaze duration data quite well. Attention-aware surprisal has strong predictability in the eye-movement data. Two types of semantic relevance significantly predicted the three eye-movement data, and the two types of enhanced metrics also performed quite significantly. In addition, the tensor interaction between *word length* and *word frequency* significantly predicts the three types of fixation duration, suggesting that both control predictors could remarkably predict eye-movements on reading. Additionally, the random effect of the *participant* is strongly significant in all GAMM fittings.

Fig. 5 displays the partial effects of these metrics on the total duration, first duration, and gaze duration in the GAMM fittings. According to Fig. 5, all surprisal metrics have a positive effect on eye-movements during multilingual reading. Conversely, the effects of semantic relevance metrics have a negative influence, implying that fixation duration decreases as semantic relevance increases. These findings suggest that language users spend more time processing words with higher surprisal values. In contrast, words with lower semantic relevance, i.e., those less likely to occur in the same context, require more time to process. Words with higher semantic relevance, on the other hand, need less time to process, as they are more likely to occur in the same context. The following presents the further indications of the performance of these metrics. With regard to semantic relevance, a lower value of semantic relevance suggests a weakened link between the target word and its context. According to memory-based theories, this smaller value indicates reduced ability to store contextual information, complicating retrieval and integration with the target word. This increases processing difficulty, requiring more time. In contrast, a larger semantic relevance implies a greater ability to store contextual representations. As a result, it needs less time to process. Moreover, a larger surprisal indicates smaller word probability, and increases processing difficulty. An increased attention-aware surprisal signifies higher surprisal values attributed to the contextual words, intensifying the processing difficulty. Moreover, while the attention-aware and enhanced approaches optimize semantic relevance and surprisal, respectively, the modified metrics still retain the fundamental characteristics of their original counterparts, as depicted in the Fig. 5.

**Figure 5:**
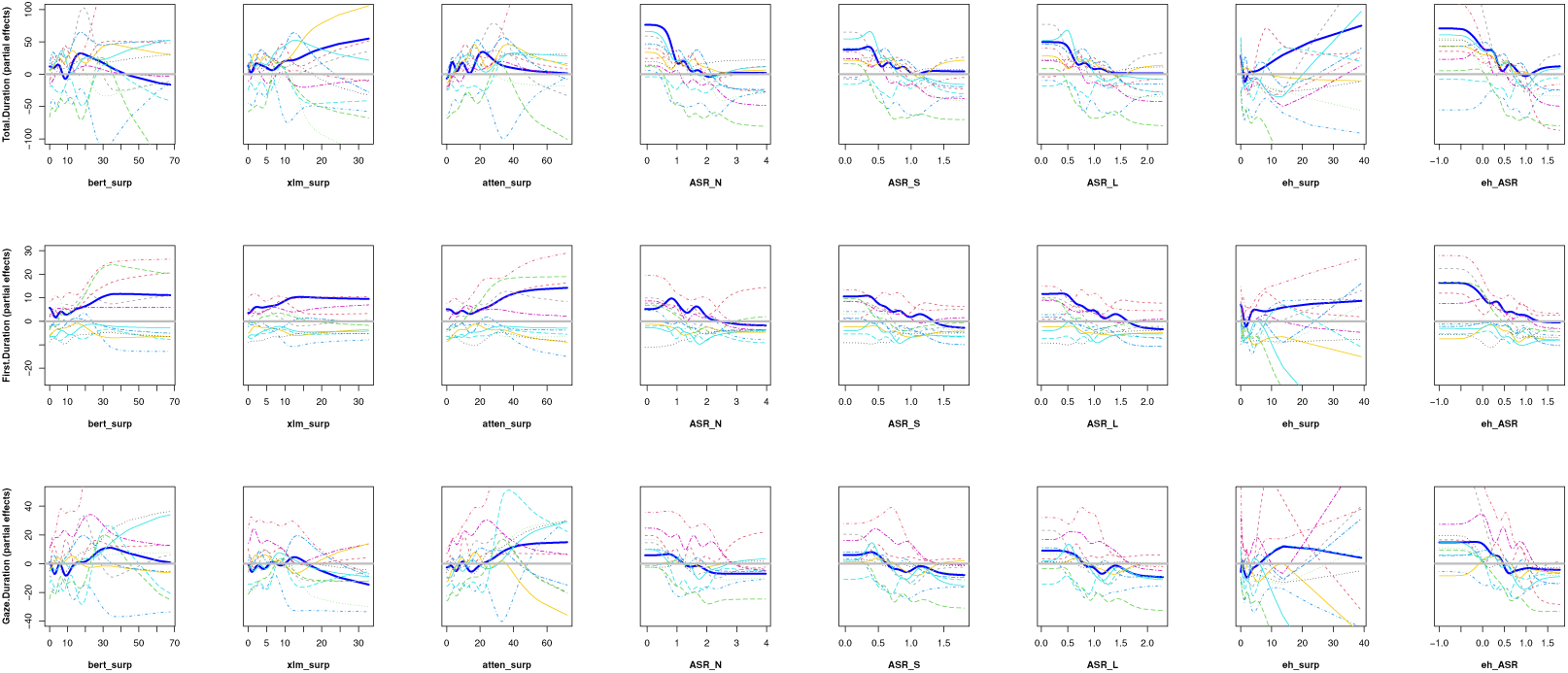
Overall partial effects of attention-aware measures and enhanced measures in prediction. The dotted lines, each in a distinct color, depict the computational metric curves for various languages, characterized by a smooth random. The solid blue curve denotes the cumulative performance for all languages.(Note: “bert_surp” = surprisal computed by m-BERT; “xlm_surp”= surprisal computed by XLM ; “atten_surp”= attention-ware surprisal; “ASR_N” = attentionaware semantic relevance (weak); “ASR_S” = attention-aware semantic relevance (small weights)); “ASR_L” = attention-aware semantic relevance (large weights));“eh_ASR”= enhanced attention semantic relevance; “eh_surp”= enhanced surprisal)

In Fig. 5, we observed that the effect of surprisal becomes relatively flat when the value of the metrics exceeds a certain threshold (e.g. 10), indicating that the surprisal effect diminishes beyond that point. However, when comparing the shape and trends of the curves, we found that the two attention-aware semantic relevance metrics (i.e., ASR_L(S), ASR_N) exhibit a much stronger and more consistent effect on the three types of duration data, in contrast to the effect of surprisal measures in Fig. 5.

To compare the performance of the metrics of our interest in these GAMM fittings, we implemented in two steps. First, we ensured that all models have a consistent number of data points and identical elements (n = 412472). And then, we verified that the variables had a significance level (*α*) with a *p*-value smaller than 0.01. Specifically, when a variable in a GAMM fitting is significant, its *p* value is smaller than 0.01. These GAMM fittings show that the control predictors, namely *word length* and *log word frequency*, are consistently significant across all cases. This suggests that sentence comprehension in naturalistic discourse reading is strongly influenced by the length and frequency of the words used. Fortunately, the metrics of interest are also highly significant in all GAMM fittings. Second, the performance evaluation of the GAMM involves comparing the difference in AIC (Akaike information criterion, i.e., ΔAIC) between a base GAMM without any proposed metrics and a full GAMM that includes one metric of our interest. This comparison helps assess the fitting performance of the GAMM, providing insights into the impact of the additional metrics on its performance. A smaller ΔAIC value indicates better model performance, while a larger value indicates the opposite. By comparing ΔAIC, we can identify which GAMM fittings are more effective and determine which metric possesses greater predictability.

As shown in Table 2, for Total Duration, we found that the ΔAIC of ASR (attention-aware semantic relevance) was much smaller than that of surprisal, and also smaller than that of atten_surp (attention-aware surprisal), but larger than that of enhanced_surp(enhanced surprisal), and also much larger than that of enhanced_ASR (enhanced attention-aware semantic relevance). enhanced_ASR consistently had the smallest ΔAIC, while surprisal had the largest value in all three cases. In comparison to surprisal, atten_surp had significantly smaller ΔAIC, indicating that it had much stronger predictability. However, attention-aware metrics of semantic relevance performed better than atten_surp. Moreover, enhanced_ASR showed the most impressive improvement in performance compared to the two attention-aware semantic relevance metrics. Additionally, the surprisal computed by m-BERT outperformed that by XLM. Taken together, these results demonstrate that the attention-aware approach in GAMM fittings leads to better performance. Furthermore, it appears that the metrics computed by the enhanced method also significantly improve their performance in GAMM fittings compared with attention-aware metrics.

**Table 2:** ΔAIC for GAMM fittings with different metrics for 13 languages (n = 407631).

| Metrics | Total Duration | First Duration | Gaze Duration |
| --- | --- | --- | --- |
| surprisal (**m**-BERT) | -939 | -244 | -663 |
| surprisal (**XLM**) | -391 | -82 | -228 |
| atten_surp | -1062 | -258 | -862 |
| ASR_N | -793 | -279 | -377 |
| ASR_S | -937 | -263 | -641 |
| ASR_L | -954 | -270 | -669 |
| enhanced_surp | -1122 | -314 | -994 |
| enhanced_ASR | -1289 | -370 | -1023 |

(“atten\_surp” = attention-aware surprisal; “ASR\_N” = attention-aware semantic relevance (no weights); “ASR\_S” = Attention-aware semantic relevance (with small weights); “ASR\_L” = Attention-aware semantic relevance (with large weights); “eh\_surp” = enhanced surprisal; “eh\_ASR” = enhanced attention-aware semantic relevance)

### 3.2 Performance for individual languages

After demonstrating the overall performance comparison, we conducted comparison for each individual language, using the same GAMM formula as previously. First, we investigated whether these metrics could significantly predict eye-movement data in each language. Second, we compared these GAMM fittings by applying the same standards.

The GAMM fitting results provide useful information about the significance of the variables included. Based on this information, we created Table 3 to show how these metrics predict eye-movements during reading in different languages. Due to the superior performance, we selected surprisal computed by m-BERT as the baseline. Consequently, we did not include the comparison results for surprisal computed by XLM. In Table 3, when the metric is not significant (we chose a threshold of *p* value being smaller than 0.01 to judge), we use ✗ to represent the case. When the GAMM fitting with the metric has the best performance, we assigned it a score of “6”. The worst performance of the GAMM fitting was rated as “1” if the metric as a variable was significant. The intermediate score will be given to other metrics. However, the *p* value in a GAMM fitting’s performance is greater than 0.001 but smaller than 0.01, we assigned it a negative symbol “(-)”. For example, when considering Dutch and total duration, surprisal does not function effectively for prediction, while “ASR_L” achieves the highest score (6) and performs the best. Following closely in the second position is “atten_surp” with a score of 5.

**Table 3:**
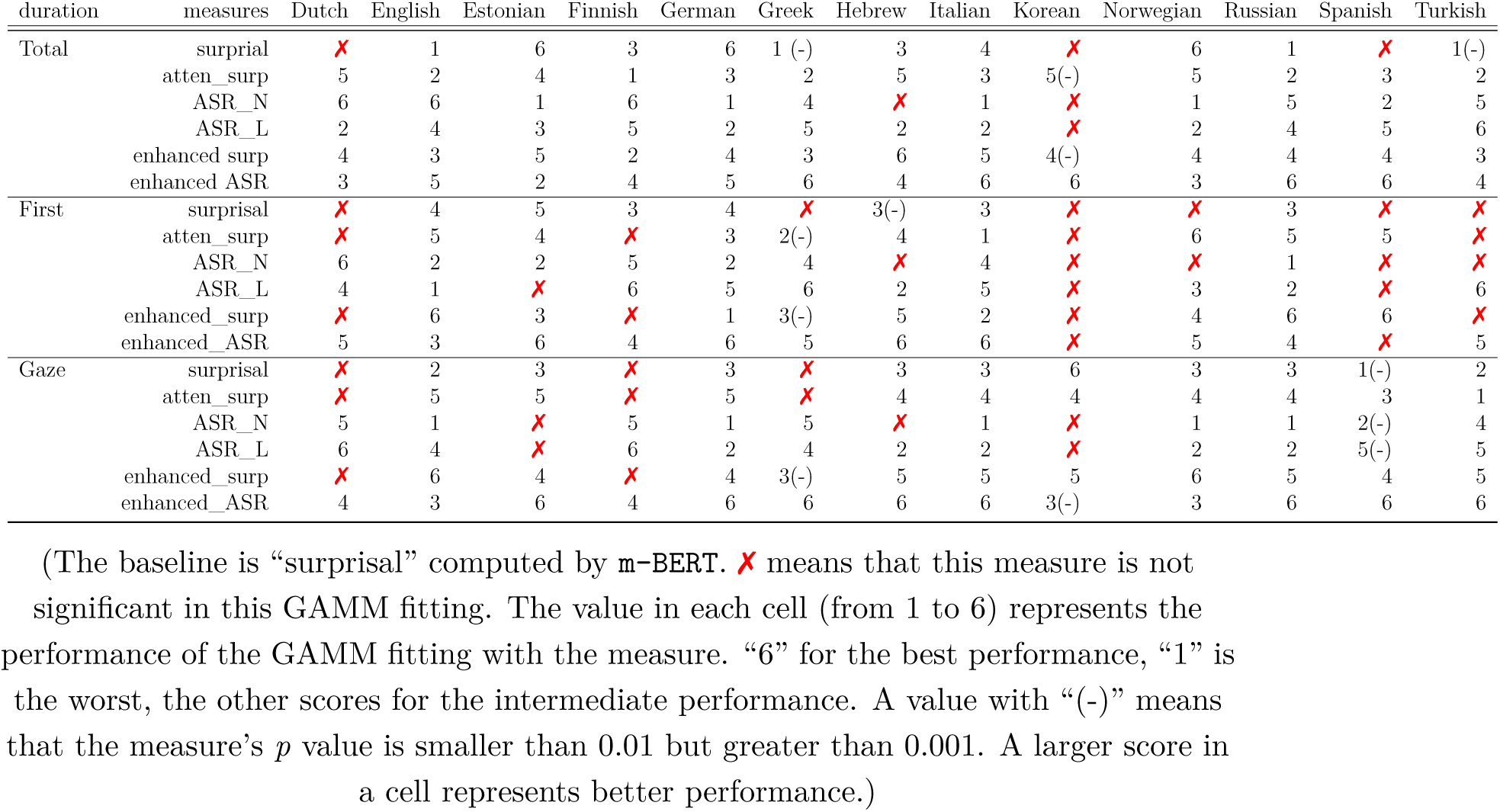
Comparison of various measures in prediction among languages (by score)

According to Table 3, surprisal was not effective in predicting Dutch, Greek, Korean, Spanish, and Turkish. Conversely, three attention-aware metrics (atten_surp, ASR_L, and ASR_N) showed superior performance compared to surprisal, which served as the baseline. However, compared with attention-aware metrics, the enhanced metrics demonstrated better prediction outcomes in terms of the number of ✗ and score, especially the enhanced_ASR, which exhibited remarkable performance by almost accurately predicting the three types of duration for all languages. This finding suggests that attention-aware metrics that consider expectation effect and weights are more effective than those that do not. Furthermore, enhanced ASR, where attention-aware information and expectation effects are combined, demonstrates significantly strong predictability. These results are basically consistent with Table 2, and the performance of these metrics in individual languages is comparable to k-fold cross-validation, confirming the effectiveness of these metrics.

The total duration is a crucial factor that indicates its overall processing difficulty, which might be more significant than the other two variables in some cases. If a metric cannot show significant predictability too often (i.e.,the number of ✗), it implies that its generalization robustness may be weakened. We also consider the total score of a given metric in each language. Three standards were taken to evaluate the performance of these metrics. Taking these factors into account along with the overall and individual performance, the measures’ ranking in terms of their effectiveness is as follows: *enhanced_ASR > ASR_L > ASR_N > enhanced_surp > atten_surp > surprisal*

Moreover, the effects of the control predictors *word length* and *word frequency* were remarkably significant, which has been reported in Kuperman et al. (2023). In addition to the effect of either predictor, both can work together to impact reading duration. The predictability of the interaction of *word length* and *word frequency* has not been reported in Kuperman et al. (2023). The details about this can be seen **Supplementary Materials (SM)**. Moreover, the information on the distribution of these metrics can be seen in **SM**.

### 3.3 Comparison of metrics computed by m-BERT and mGPT

Due to the fact that mGPT cannot process the Norwegian language, we had to report the relevant results using mGPT in this separate section. The Norwegian dataset in the MECO was excluded in the testing dataset of the MECO (12 languages). The attention-aware surprisal was based on surprisal computed by mGPT, and enhanced metrics were also estimated using mGPT-based surprsial. Adopting the same evaluation standards, we created Table 4 to show the result of comparison of GAMM fittings.

**Table 4:** ΔAIC for GAMM fittings with different metrics for 12 languages (n = 329728).

| Metrics | Total Duration | First Duration | Gaze Duration |
| --- | --- | --- | --- |
| surprisal (<u>m</u>-BERT) | -385 | -155 | -362 |
| surprisal (<u>XLM</u>) | -318 | -66 | -179 |
| surprisal (<u>m</u>GPT) | -482 | -289 | -403 |
| atten_surp | -695 | -70 | -438 |
| ASR_N | -546 | -311 | -448 |
| ASR_S | -653 | -451 | -319 |
| ASR_L | -323 | -329 | -457 |
| enhanced surp | -1007 | -337 | -473 |
| enhanced ASR | -1121 | -351 | -521 |

“atten\_surp” = attention-aware surprisal based on the surprisal computed by m-BERT;
“ASR\_N” = attention-aware semantic relevance (no weights); “ASR\_S” =
Attention-aware semantic relevance (with small weights); “ASR\_L” = Attention-aware semantic relevance (with large weights); “eh\_surp” = enhanced surprisal; “eh\_ASR” =
enhanced attention-aware semantic relevance

Following the standards of comparison mentioned above, we found that the predictability of surprisal computed by mGPT was slightly stronger than that computed by m-BERT. Unfortunately, the surprisal computed based on XLM exhibited the poorest performance. The three types of word surprisal computed by different LLMs did not surpass the predictive ability of the attention-aware and enhanced metrics proposed in the current study for predicting eye-movements on reading. The other metrics have the similar performance shown in Table 4. The results in Table 4 are basically consistent with Table 2. Following the methods in Table 3, we conducted comparison for each individual language (without Norwegian), using the same GAMM fittings as before. The result is basically consistent with Table 3, and it is seen in **SM**.

Through comparisons at different levels, we found that the results are generally consistent. Overall, the performance of these metrics can be ranked as : enhanced metrics > attention-aware semantic relevance > attention-aware surprisal > surprisal .

## 4 Discussion

In summary, the GAMM analyses consistently produced results that revealed the following: first, it became evident that factors such as *word length* and *word frequency* yield substantial influence over human reading behavior across various languages. Second, the metrics generated by the approaches we proposed have a pronounced impact on human reading behavior. This highlights the intertwined impact of both expectation and memory effects on shaping reading behavior.

The following summarizes the predictability of the metrics we adopted on the eye-movement on reading. First, word *surprisal* was a strong predictor for all three response variables in overall performance and across a number of languages, which is consistent with numerous studies on word predictability effect and surprisal effect on reading and language processing (Staub, 2015; Hale et al., 2015). Second, both attention-aware metrics and enhanced metrics emerged as stronger predictors for all three response variables, demonstrating superior performance across the majority of languages in the MECO dataset, surpassing the effectiveness of *surprisal*. Third, the enhanced metrics were superior to the attention-aware ones. After comparing the overall and individual results for each language, we found that the enhanced attention-aware semantic relevance (i.e., enhanced ASR) consistently performed the best. The attention-aware semantic relevance also outperformed the attention-aware surprisal in predicting fixation durations. Forth, a significant finding was the reinforcement of the preview effect in naturalistic reading, further emphasizing its role in guiding and improving the reading process.

Further, the achievements of these metrics strongly underscore the substantial influence of contextual information on language processing. For instance, the remarkable predictability of attention-aware semantic relevance highlights a crucial insight: when readers process words in context, they are significantly influenced by its semantic connection within the local context. Additionally, the improved performance of attention-aware surprisal further reinforces the notion that readers are indeed influenced by the surprisal of contextual words. As readers process a text, the level of unexpectedness or surprisal associated with the surrounding words significantly impacts their comprehension and interpretation of the content. Attention-aware metrics offer a more comprehensive understanding of how readers navigate through text and adapt their cognitive resources. Further, the resounding success of enhanced metrics underscores how integrating diverse information sources can greatly enhance the simulation of realistic cognitive reading mechanisms. A more detailed analysis of these metrics is provided in the following sections.

### Memory-based strengths and parallel processing

The attention-aware approach is highly capable of incorporating contextual information, and is intrinsically linked to memory, operating on principles of storage, decay, and retrieval of contextual information. The following details how the attention-aware approach realized storage, and decay on contextual information based on memory, and then discusses why attention-aware metrics can simulate various reading theories/language processing models, and quantitatively evaluate them.

The preceding words prior to the target word in a window, as illustrated in Fig. 2A, B and Fig. 6, simulate a short-term memory stack, mirroring how readers retain memory of previously encountered words and their meanings. The method for using various weights of semantic relevance between any two words is inspired by both the attention mechanism found in Transformers and the human process of forgetting. The weight values gradually decrease with the distance between the target word and the contextual word, similar to the forgetting curve (Murre and Dros, 2015). Words that are closer to the target word are given more weights, reflecting their greater relevance and likely impact on comprehension and memory. The positional distance between words can in our model be thought of as the time interval between learning and recall in memory models like the ACT-R model (Anderson et al., 1997). Both the stack of words and the forgetting curve involve a form of memory “decay” or decrease with the time, that is, weights decreases as positional distance in the stack increases, and memory retention decreases as time distance (interval) increases. Fig. 3A shows the similarity between the forgetting curve and word positional weights in the attention-aware approach. Additionally, different weights can also help encode the information of word order in the memory stack. In short, this could be linked to memory models in terms of how memory is decayed during the encoding of information, which subsequently affects how humans process words during reading. More importantly, the attention-aware approach is computational and fundamentally memory-based, facilitating memory storage, retrieval, and integration. This approach not only realizes the memory function but also incorporates the expectation effect, as depicted in the Fig 6.

**Figure 6:**
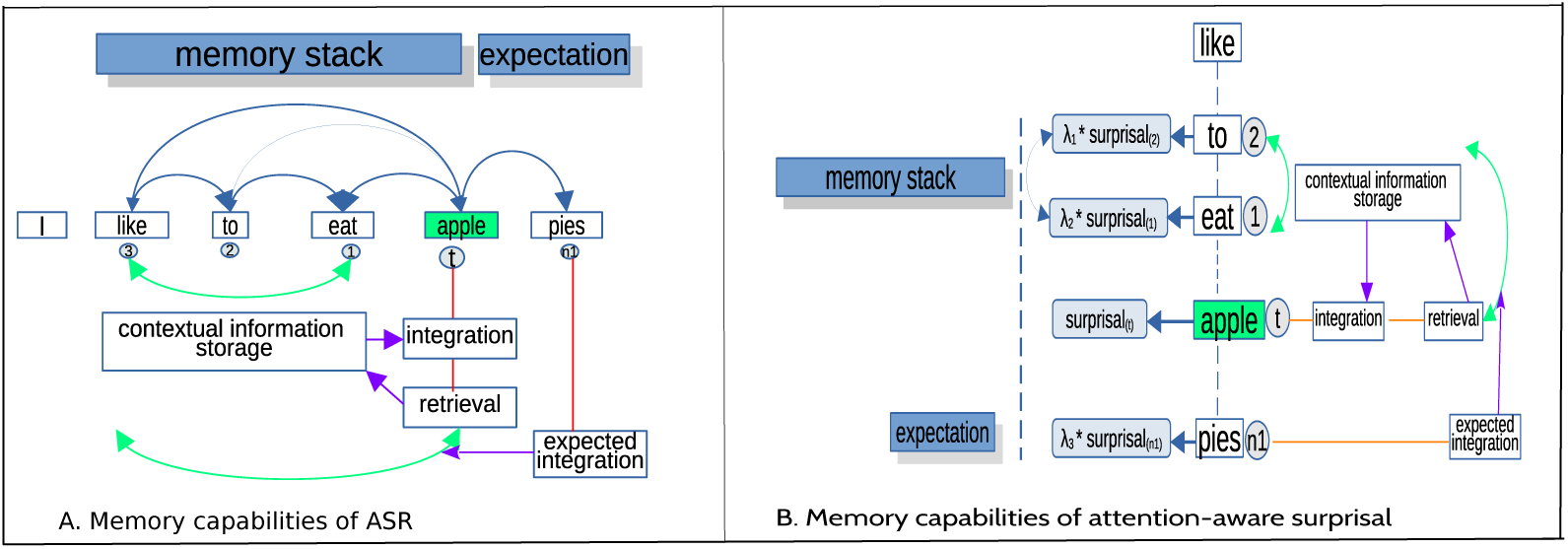
Memory capabilities of attention-aware approach. (In Panel A, attention-aware semantic relevance is showcased, demonstrating memory storage, retrieval of the storage, the integration of the target word, as well as the expected integration of “n+1” word. Meanwhile, Panel B portrays attention-aware surprisal, also exemplifying memory storage, retrieval, and integration.)

Our results provide empirical evidence to support this viewpoint. The attention-aware metric without using weights, denoted as ‘ASR_N’ did not surpass the metric that incorporates weights, denoted as “ASR_L(S)” in predicting eye-movements, shown in Tables 2 and 4. The metric without using weights is nearly equivalent to not considering the forgetting impact on reading, leading to inferior performance compared to the metric that considers weights. This finding suggests that the metric that considers the human forgetting mechanism and word order information exhibits stronger predictability than the one that disregards this factor.

Our results also reveal the importance of short-term memory in reading and language processing. Previous research on language comprehension models has allocated minimal attention to working memory, with notable exceptions being Baddeley (2000) and Gibson (1998). When contrasted with other established memory-based metrics, such as dependency locality theory (Gibson, 1998), which falls short in predicting eye-movements during English reading (Demberg and Keller, 2008; Kwon and Sturt, 2014), our proposed semantic relevance metrics which are short-term memory based exhibit a robust capacity to predict eye-movements in multilingual reading. Moreover, debates over the nature of working memory continue to persist. Some scholars, such as Baddeley (1992), argue that fleeting memories are housed in a specialized temporary storage area. Others, like Buchsbaum and D’Esposito (2019), believe that these memories arise from language comprehension and various cognitive functions. The challenge in resolving these debates has been exacerbated by a lack of quantitative studies on how transient memory emerges from language processes (Norris, 2017). Many existing computational models that explore emergent working memory are narrow in focus, mainly concentrating on how individual words are retained (Ueno et al., 2011). Our work takes a different approach, introducing semantic relevance metrics that provide a broad computational framework. This framework not only illuminates the nature of short-term memory in the context of understanding sentences and discourse but also examines verbal working memory as a phenomenon that naturally arises from the process of comprehending language (Schwering and MacDonald, 2020).

Next, we focus on how the proposed metrics contribute to understanding various theories and models on reading and language processing. The debates on reading behavior and paradigm have sparked significant interest in language processing (Clifton Jr et al., 2016; Reichle et al., 2003; Schotter et al., 2012; López-Peréz et al., 2016; Brothers et al., 2017; Li et al., 2022). For over two decades, two prominent models, the E-Z Reader and the SWIFT model, have remained at the center of heated discussions within extensive research endeavors(Engbert et al., 2005; Pollatsek et al., 2006). The E-Z Reader suggests a sequential word recognition process, similar to processing letters in an alphabetic sequence (i.e., serial processing). In contrast, the SWIFT model proposes a more flexible approach, integrating information from a broader visual span and utilizing contextual cues to facilitate word recognition, potentially allowing readers to skip certain words during fixations for a more efficient reading process (i.e. parallel processing). During SWIFT reading, a reader often makes a regressive saccade, a backward eye movement, to reinspect a word or a portion of the text. Specifically, the processing of the target word can be influenced by upcoming words (“n+1” word, or “n+2” word in some studies), regardless of whether our eyes are currently fixated on them. This is referred to as “parafoveal processing”. Essentially, during reading, parafoveal information can impact the processing of the currently fixated word (parafovea-on-fovea effect, or previewed benefit), and words perceived parafoveally can facilitate their subsequent processing. However, the way word order is encoded presents a significant challenge in the SWIFT model used in sentence reading. Understanding the sequence of words is essential for comprehension Fig. 3 B(a),(b) illustrate preview reading, serial and parallel processing. Such effects have been confirmed by numerous empirical studies (Kennedy and Pynte, 2005; López-Peréz et al., 2016; Vasilev and Angele, 2017). These effects have also been supported by neural evidence (Niefind and Dimigen, 2016; Pan et al., 2021).

Our attention-aware approach enables effortless simulation of both E-Z reader and SWIFT model. It generated several metrics, with the first (“ASR_N”) resembling E-Z reader and the latter two (“ASR_L” (or “ASR_S”) and attention-aware surprisal) simulating the SWIFT model. Surprisal, a metric that only considers the preceding context, regrettably does not account for the upcoming context, so it can simulate E-Z reader. In contrast, the attention-aware surprisal considering the information of the next word’s surprisal can simulate the SWIFT model. The “ASR” incorporates information from the next word, simulating the SWIFT model. The word order information can be encoded by our metrics, effectively addressing the issue present in the SWIFT model. In contrast, “ASR_N” simulates E-Z reader without including the next word information. Due to the combination of attention-aware surprisal and attention-aware semantic relevance, the enhanced metrics also include information from the next word and word order. Therefore, the enhanced metrics can similarly simulate the SWIFT model. Fig. 3B(c) displays the simulation of our metrics for different language processing models, and their results of their quantitative evaluations.

The comparison of the metrics through GAMM analysis allows for the quantitative evaluation of these theories or models. Specifically, the superior performance of “ASR_L(S)” than “ASR_N” support the next-word benefit. The attention-aware metrics also surpassed surprisal in prediction, further highlighting the benefit of considering next-word information. Attention-aware surprisal, encompassing information about the “n+1” word, demonstrated superior performance compared to surprisal alone (excluding n+1” information). This enhancement also aligns with the validation of preview benefits. The influence of the predictability of the “n+1” word on the target word has been substantiated by numerous studies (Kliegl et al., 2004; Kennedy et al., 2013; Huck et al., 2017), and our study is consistent with the findings in these studies. Generally, the metrics considering the following word information outperformed the ones without incorporating the information of the next word. It validates the existence of preview benefits in massive multilingual eye-movement datasets, further supporting the SWIFT model and parallel processing.

However, the results show that ‘ASR_N’ could slightly outperform ‘ASR_L(S)’ incorporating the information of “n+1” word in predicting first fixation duration. This could be attributed to the fact that the initial stage of word processing (represented by first duration) is significantly influenced by the left context. Preview effects, on the other hand, are likely to impact word processing at a later stage. This could explain why ‘ASR_N’ did not surpass the other attention-aware semantic relevance metrics in predicting total fixation duration and gaze duration. Despite this, the performance of ‘ASR_N’ is influenced by the particular languages and types of fixation durations involved. For instance, ‘ASR_N’ appears to outperform ‘ASR_L’ in predicting total durations for English, Dutch, Finnish, and Russian. However, in the case of gaze durations, ‘ASR_L(S)’ outperformed ‘ASR_N’ in the majority of languages. All this suggests that the advantage of using preview information is not necessarily significant across all languages. Taken together, attention-aware metrics offer the great potential to conduct comprehensive simulations and quantitative assessments across various reading paradigms and language processing models.

Some lab-based experiments on reading carefully controlled sentences seem not to support parafoveal effects. In contrast, reading in labs without control is closer to naturalistic discourse reading rather than lab-controlled sentence reading. Our results show that the effects of surprisal and attention-aware metric without the next word information do occur in eye-movements, supporting the existence of E-Z models on reading and serial processing in language comprehension and processing. However, the stronger predictability of attention-aware metrics considering the next word information suggests that SWIFT model is prominent in naturalistic discourse reading and parallel processing could easily take place during realistic reading. However, when reading takes place under controlled conditions (such as reading controlled sentences in labs), serial processing might overshadow parallel processing. This could explain why both phenomena can be observed and both can be inter-activated with each other, but the prominence of each depends on the specific conditions (Wen et al., 2019; Jensen et al., 2021) and different languages.

### The advantages of enhanced approach

In the previous section, we explored the strengths of the attention-aware method. Our analysis shows that the enhanced metrics we used were superior in performance. Now, we delve into the specific advantages of the enhanced approach.

First, the enhanced approach incorporates both memory and expectation information, providing a more comprehensive representation of how humans comprehend and process language in real-world contexts. Humans rely on both their memory of previously encountered words and their expectations of upcoming words to make predictions and efficiently comprehend the meaning of a sentence. The expectation and memory should be mutually interacted (Ryskin and Nieuwland, 2023) in language comprehension and processing. Our enhanced metrics could quantitatively evaluate these viewpoints. When the model considers both memory-based contextual semantic relevance and expected-based surprisal, it better reflects and simulate the dynamic interplay between memory updating and expectation adjustment during reading. The metrics generated by such a model become more precise in capturing how readers allocate their attention, make eye movements, and adapt their reading strategies based on the meaningfulness of the text and the degree of unexpectedness they encounter. In the current study, we calculated enhanced metrics from two perspectives: (1) semantic relevance dominated with weighted surprisal (“enhanced ASR”), and (2) surprisal dominated with weighted semantic relevance (“enhanced surp”). For example, when surprisal is incorporated into the memory-based model, the enhanced metric (“enhanced ASR”) can better capture moments of increased cognitive processing due to unexpected words or phrases. This provides a more accurate representation of the dynamic and adaptive nature of the reading process, as readers adjust their reading strategies based on the unexpectedness of the text. On the other hand, after semantic relevance is incorporated in surprisal, the enhanced metric (“enhanced surp”) can better account for the facilitation or disruption caused by the meaningfulness and coherence of the text. Note that enhanced ASR basically aligns with semantic relevance in predicting eye-movement, while enhanced surprisal maintains similar attributes to standard surprisal for predicting eye-movements, as illustrated in Figure 5.

Second, the enhanced approach under discussion can be characterized as a method of weighted superposition, a concept that has found extensive application in both quantum physics and engineering disciplines (Dirac, 1981). Superposition, in a broad sense, pertains to the phenomenon wherein, for a given linear differential equation, the aggregate of two solutions emerges as a solution in itself, such as wave interference (Fig. 4A). Further, the superposition method, along with other quantum techniques, has been extensively employed in cognitive sciences, including areas such as memory, decisionmaking, and categorization (e.g., Bruza et al., 2015; Busemeyer and Wang, 2015; Pothos and Busemeyer, 2022).

The enhanced approach under consideration draws inspiration from the principle of superposition and quantum cognition. The reading state may be conceptualized as a superposition state from the perspective of the quantum cognition, wherein various factors retain the potential for expression. Certain factors may interfere with one another, contributing constructively and ultimately altering the final observed measurements, such as eye-movement measures recorded by an eye-tracker (i.e. fixation duration, saccades etc.). This explains why ‘ASR’ and ‘attention-aware surprisal’ may be superposed to create constructive interference. However, weights should be employed to adjust the value of subordinated metric when an effect is perceived as dominant, given the highly adaptable reading strategies humans take. In the context of cognitive science, the superposition collapse state is not inherently subject to instantaneous measurement or recording due to its potential characterization as a cognitive operation (Fig. 4B). Nonetheless, recourse to statistical analysis affords an avenue through which to elucidate the influence exerted by such superposition states on observable indicators. In this way, by integrating both types of information through the weighted superposition (Fig. 4C), the enhanced metrics offer a more nuanced simulation of human language processing from the perspective of quantum state, thereby contributing to a more precise comprehension of cognitive processes (Fig. 3B(c)).

Third, the enhanced metrics are more easily computed. As mentioned earlier, the lossy-context theory (Futrell et al., 2020; Hahn et al., 2022) once merged dependency lengthy information (dependency locality theory from Gibson (1998)) into surprisal and created lossy-context surprisal. However, the context-lossy surprisal theory lacks a fully implemented specification regarding the specific elements of preceding context that are susceptible to memory loss. This shows that the lossy-context theory has lacked a feasible computational implementation and limits its broad applicability for comprehensive quantitative evaluations of word processing. Surprisal has also demonstrated shortcomings in various languages, with its predictability falling short when compared to its semantic relevance metric counterparts. This reveals that the effect of expectation (or probabilistic inference) is not as robust as previously anticipated. In essence, the theory of lossy-context surprisal contends that the predominant factor is the effect of expectation (surprisal), albeit modulated by memory information (Hahn et al., 2022). This perspective aligns with a serial processing framework. In contrast, our approach posits a simultaneous occurrence of expectation and semantic plausibility effects in contexts (i.e., parallel processing plus word order), even though these two influences can mutually affect each other. The concept of a superposition state involving the two previously mentioned effects also is also in line with parallel processing. Compared with lossy-context surprisal, it is evident that the enhanced approach solidifies the distinct contextual boundaries, thereby enabling the practical execution of computations for metric generation. In short, our method offers easy computation to generate the metrics incorporating both memory and expectation information, and the most significant advantage lies in their strong predictability for almost all words in the corpus. The approach allows us to compute metrics and validate their predictability conveniently and effectively.

### Main methodological contributions

This section offers a comprehensive analysis of the key methodological contributions in the current study. This study’s **primary** contribution is the advancement of our computational methodology. First, we proposed two approaches to optimize existing metrics and generated a number of metrics. We introduced an attention-aware approach to create a number of metrics. The attention-aware approach is not confined to optimizing surprisal or semantic relevance; it can also be used to improve other metrics. We further created an enhanced approach to create more powerful metrics. Second, these metrics showed their superiority in predicting and explaining language processing compared to the existing metrics. Third, our approaches and metrics are highly interpretable from a cognitive perspective. Forth, most cognitive models exhibit computational limitations, while a significant portion of computational models lacks interpretability. The availability of explainable computational models for cognition remains notably limited. The potential for our computational methods and metrics to foster interpretable computational models in cognition is significant.

The current study’s **second** contribution highlights the usefulness of computational metrics in predicting language processing and comprehension across multiple languages, not only English. Our findings support previous research that demonstrates the predictability of contextual semantic relevance in eye movements during reading and comprehension (e.g., Roland et al., 2012; Frank and Willems, 2017; Sun et al., 2023a). We also found that surprisal effects were present in multiple language processing, which actually supports numerous studies using surprisal to predict language processing (Hale et al., 2022; Hale, 2016; Schrimpf et al., 2021). The results show that our new metrics have generalization across languages (differ from language families, script types, morphological typology, and orthographic transparency, see Table 1 in Siegelman et al. (2022)), but performed differently in terms of language and context (Li et al., 2022). Unifying the generalization of cognitive phenomena across languages is one important goal of computational cognitive science. These metrics can perform complex tasks and to handle large amounts of data on language processing.

The **third** contribution is that our attention-aware and enhanced approaches introduced a set of convenient metrics for the quantitative evaluation of various reading theories and language processing models, as previously discussed. Specifically, we have applied these metrics to compare the E-Z Reader and SWIFT models (Pollatsek et al., 2006; Engbert et al., 2005), representing serial and parallel processing (Snell and Grainger, 2019), respectively. These models have been validated through lab-based experiments and limited datasets, but the scalability of our approaches allows for more extensive testing with massive multilingual datasets. Our approaches can efficiently process and analyze large datasets, enabling researchers to assess these theories using real-world data from diverse reading scenarios. For instance, E-Z Reader and SWIFT models assume that some factors, such as word frequency (a memory-based measure), semantic plausibility, and word predictability (an expectation-based measure), are significant during reading. However, these models heavily rely on these external factors to estimate their performance and lack the capacity to compute their own metrics for evaluating and predicting eye-movements. In contrast, the attention-aware and enhanced approaches we proposed are capable of independently generating a variety of metrics. These metrics are able to evaluate eye-movements, as well as other data related to language comprehension.

### Extensions

The approaches and metrics we have introduced can likewise be effectively applied to predict eye-movements on reading in more languages, including Chinese, Danish, and Portuguese. Our another study demonstrates the applicability of these metrics in predicting phonetic and acoustic information in spontaneous speech across multiple languages, including speech duration, intonation, pitch rate, and more. Spontaneous speech represents the phenomenon of language production. Interestingly, the metrics that incorporate next-word information perform even better in predicting phonetic features in spontaneous speech. This indicates the potential applicability of our approaches and metrics in language production research.Moreover, the metrics proposed can predict and explain neural data on naturalistic discourse processing (e.g., EEG and fMRI signals, EEG aptitude change rate from 300ms to 500ms). Such metrics can be also used to predict how humans process visual information (e.g., fixation numbers in objects in an image). In other words, such methods and metrics can be applied to interpret and explain how humans process multi-modal information. Unification of theories and models can further help to provide a more complete understanding of human cognitive and neural mechanisms on processing multi-modal information.

In the future, we can apply these methods to predict neural activities in language processing for a wide range of languages using existing multilingual neural datasets (e.g. fMRI BOLD datasets (Malik-Moraleda et al., 2022)). Additionally, by employing a quantum-inspired framework, it becomes possible to capture the intricate and multifaceted nature of human language processing. In forthcoming endeavors, the framework and algorithms rooted in quantum cognition could also be harnessed to potentially yield profound insights into the underlying mechanisms governing human comprehension processes.

To sum up, the present study presents compelling evidence supporting working memory advantages, semantic preview benefits and parallel processing in multilingual reading. These findings unquestionably contribute to resolving the contentious debates surrounding human reading and language processing. The metrics proposed in this study are highly interpretable and contribute to a better understanding of serial or parallel processing in language comprehension. This makes them valuable computational tools for modeling language comprehension, production and processing across languages. The insight conversely helps design more powerful algorithm in deep learning and AI.

## The Data Availability Statement

The training dataset is available at https://fasttext.cc/docs/en/crawl-vectors. html. The BERT and GPT models are available at https://huggingface.co/bert-base-multilingual-cased and https://huggingface.co/ai-forever/ mGPT respectively. The testing dataset is available at https://osf.io/ 5fygd/?view_only=4ac60ab05ec04cef8c14c3c19b26efaa. The code used in the present study is available at https://osf.io/ghsq4/. All code used to compute surprisal and attention-aware semantic relevance is available at: https://osf.io/5fygd/.

## Authors’ Contributions

K. S. conceived of the presented idea. K.S. developed the theory and performed the computations. K.S. wrote and edited the manuscript.

## Funding

No funding was obtained for this study.

## Conflict of Interest Statement

The authors have no conflicts of interest to declare.

## A Correlations among the metrics

The Pearson correlations among various metrics utilized in the current study are illustrated in Fig. 7. The *ρ*-value for each correlation is less than 0.01, indicating significance. Specifically, the correlation coefficients between surprisal metrics (bert, xlm, gpt) and ASR metrics (across three variants) range from -0.04 to -0.09. Additionally, ASR metrics exhibit low correlation with attentional surprisal and enhanced surprisal, with coefficients ranging from -0.03 to -0.13. These findings suggest that ASR and surprisal metrics are distinct and measure different constructs.

**Figure 7:**
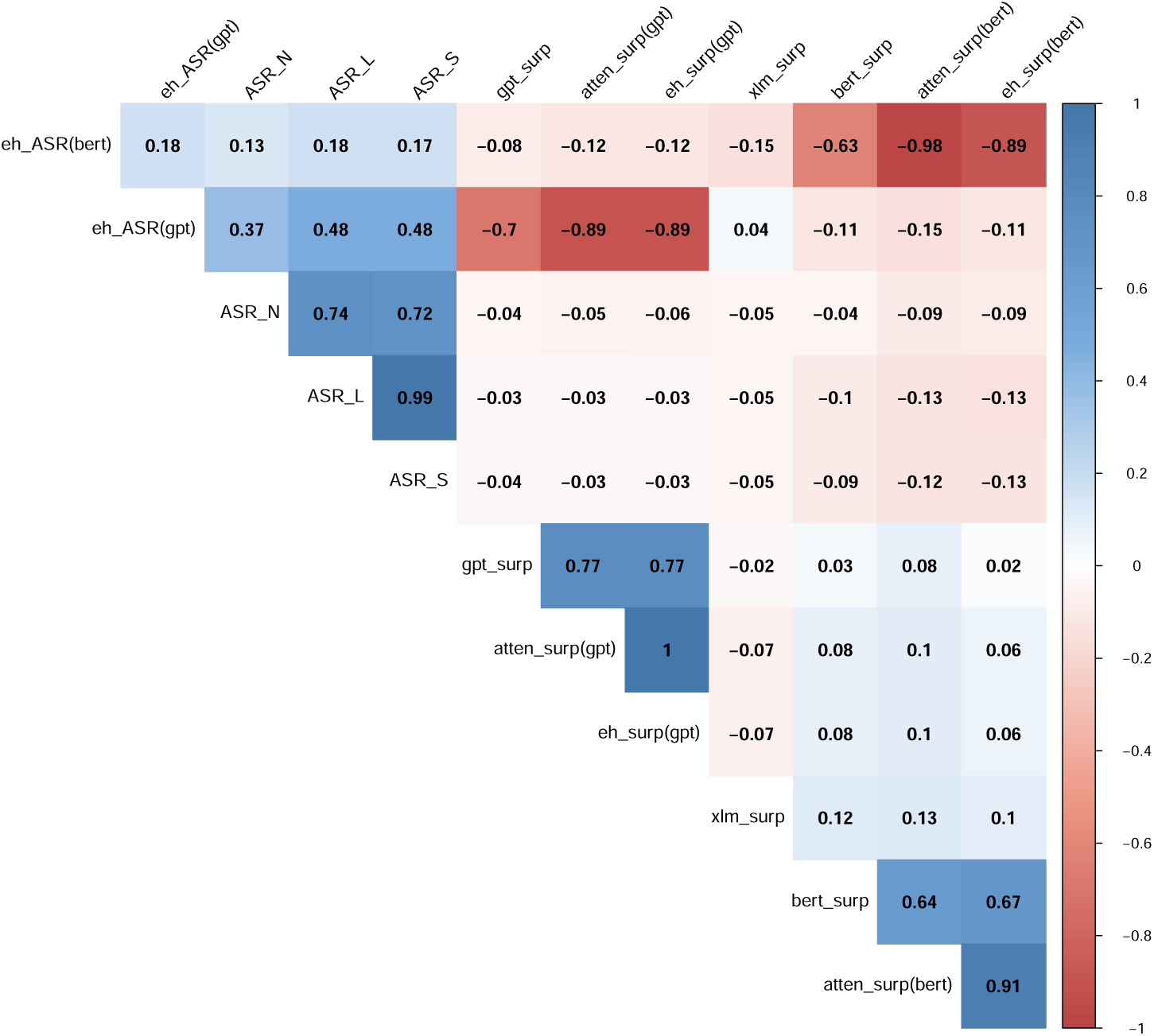
Pearson correlations among various metrics in the present study.(Note: “bert_surp” = surprisal computed by m-BERT; “xlm_surp”= surprisal computed by XLM ; “atten_surp”= attention-ware surprisal; “ASR_N” = attention-aware semantic relevance (weak); “ASR_S” = attention-aware semantic relevance (small weights)); “ASR_L” = attention-aware semantic relevance (large weights));“eh_ASR”= enhanced attention semantic relevance; “eh_surp”= enhanced surprisal)

**Figure 8:**
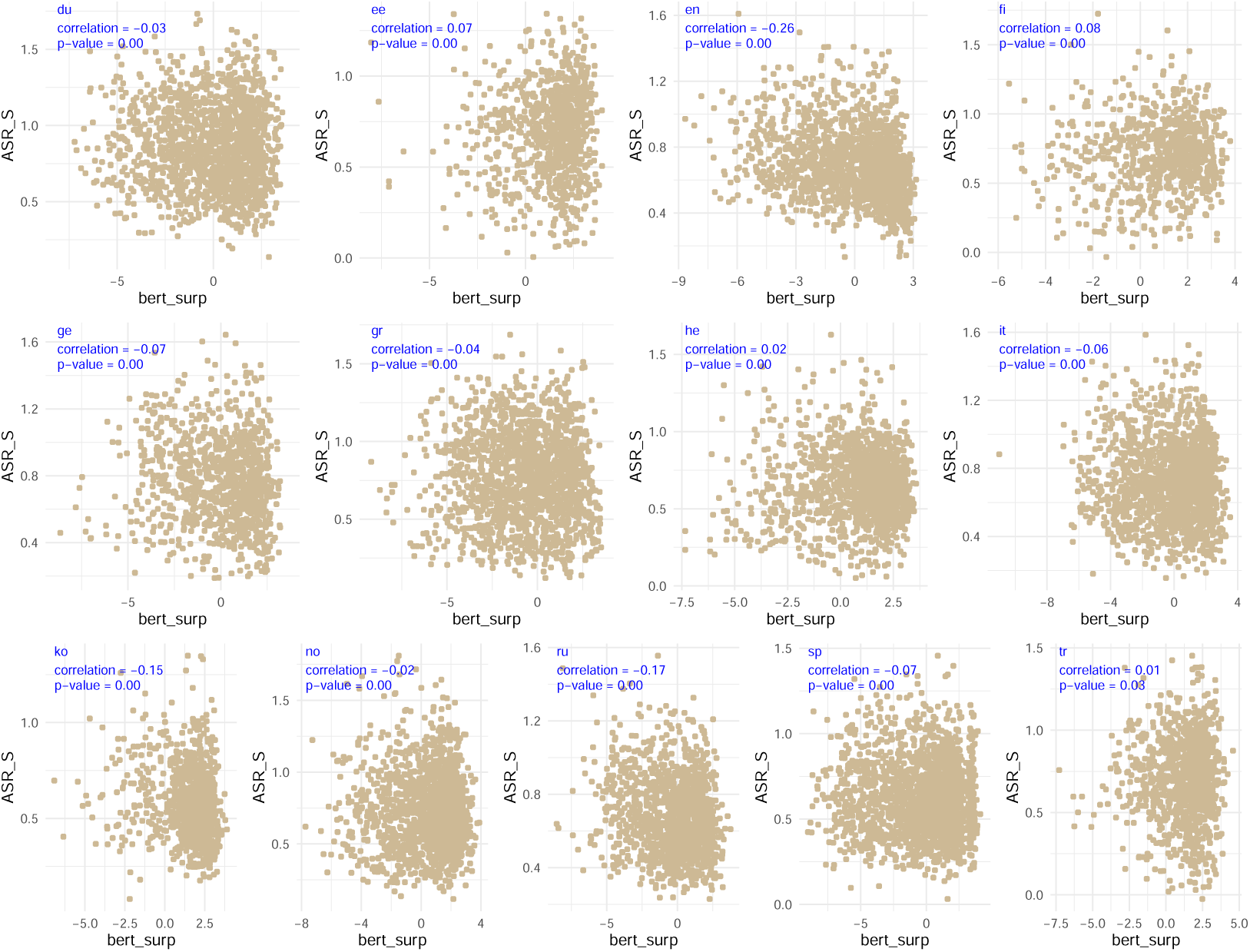
Correlations between surprisal computed by BERT and ASR in each language.(Note: “bert_surp” = surprisal computed by m-BERT; “ASR_S” = attention-aware semantic relevance (small weights)); du = Dutch; en = English; ee = Estonian; ge = German; gr = Greek; fi = Finnish; he = Hebrew; it = Italian; no = Norwegian; ko = Korean; ru = Russian; sp = Spanish; tr = Turkish within each language. Given the significant disparity between surprisal values derived from BERT and GPT, we opted to assess their correlations with ASR_S independently across languages. Fig. 8 illustrates the correlation between surprisal computed by BERT and ASR_S, whereas Fig. 9 depicts the correlation between surprisal computed by GPT and ASR_S. Across all examined languages, we observed that the correlation between surprisal metrics and ASR is uniformly weak. This suggests that the two metrics are distinct within each language context. These findings align with those presented in Fig. 7, further substantiating the notion that surprisal and ASR metrics capture different aspects in language.

**Figure 9:**
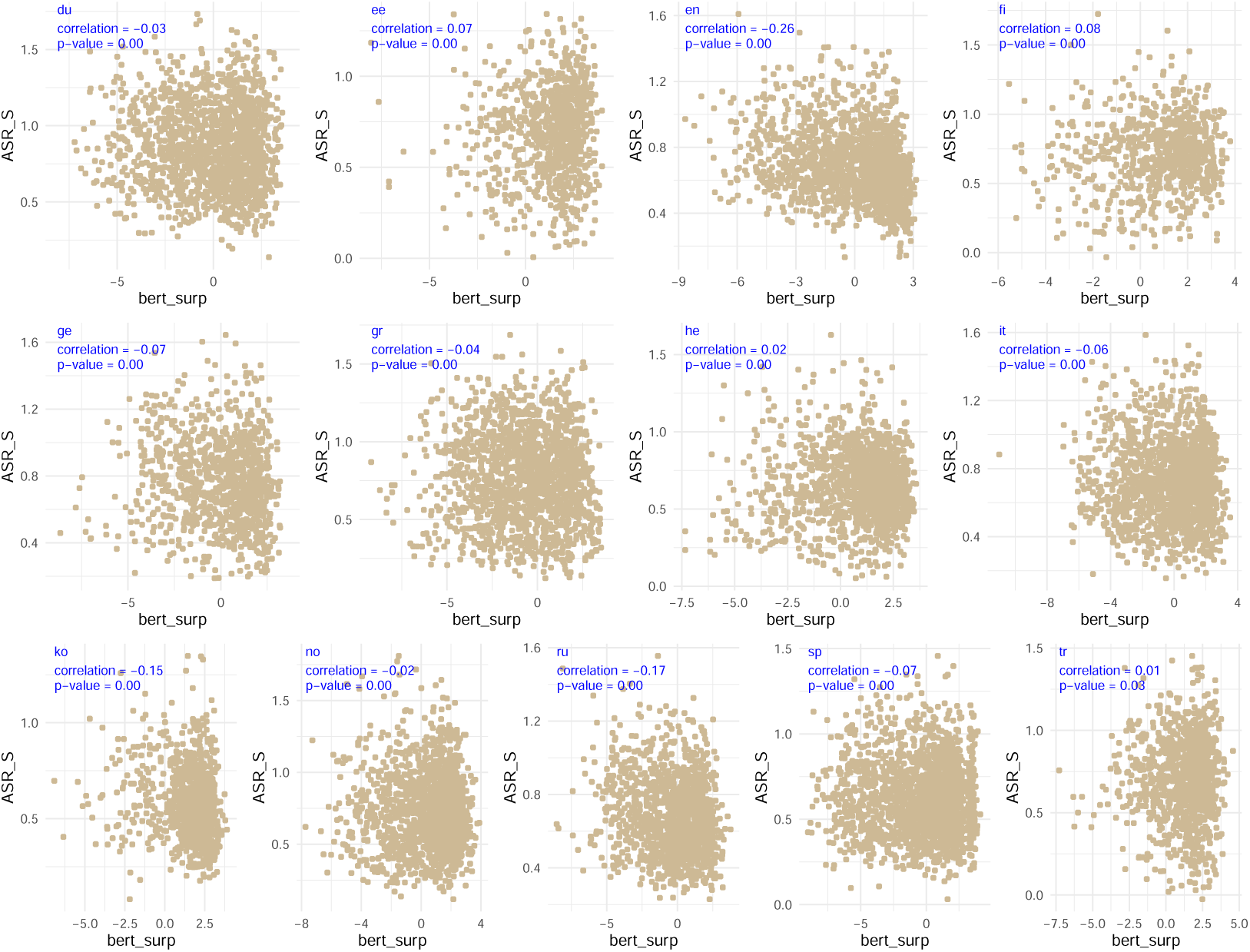
Correlations between surprisal computed by GPT and ASR in each language.(Note: “gpt_surp” = surprisal computed by mGPT; “ASR_S” = attention-aware semantic relevance (small weights)); du = Dutch; en = English; ee = Estonian; ge = German; gr = Greek; fi = Finnish; he = Hebrew; it = Italian; no = Norwegian; ko = Korean; ru = Russian; sp = Spanish; tr = Turkish

The relationship between ASR metrics and enhanced ASR metrics presents a more nuanced picture. ASR_S demonstrates a weak correlation with enhanced_ASR (bert) but a strong correlation with enhanced_ASR (bert). Furthermore, surprisal computed by BERT shows a weak correlation with surprisal computed by GPT. Attentional surprisal and enhanced surprisal based on BERT also display weak correlations with their counterparts based on GPT. However, attentional surprisal and enhanced surprisal based on BERT are strongly correlated with surprisal metrics computed by BERT, a pattern that holds true for GPT-based metrics as well.

This section elucidates the correlation between surprisal and ASR metrics across different languages. Specifically, we analyzed surprisal values computed by BERT and GPT, alongside ASR_S, to examine their correlations

## Notes

### Competing Interest Statement

The authors have declared no competing interest.

### Summary of Updates

Add new information on correlations among the metics.

## References

1. Anderson, J. R., Matessa, M., and Lebiere, C. (1997). Act-r: A theory of higher level cognition and its relation to visual attention. Human– Computer Interaction, 12(4):439–462.

2. Armeni, K., Willems, R. M., and Frank, S. L. (2017). Probabilistic language models in cognitive neuroscience: Promises and pitfalls. Neuroscience & Biobehavioral Reviews, 83:579–588.

3. Baddeley, A. (1992). Working memory. Science, 255(5044):556–559.

4. Baddeley, A. (2010). Working memory. Current Biology, 20(4):136–140.

5. Baddeley, A. D. (2000). Short-term and working memory. The Oxford Handbook of Memory, 4:77–92.

6. Blasi, D. E., Henrich, J., Adamou, E., Kemmerer, D., and Majid, A. (2022). Over-reliance on english hinders cognitive science. Trends in Cognitive Sciences.

7. Brothers, T., Hoversten, L. J., and Traxler, M. J. (2017). Looking back on reading ahead: No evidence for lexical parafoveal-on-foveal effects. Journal of Memory and Language, 96:9–22.

8. Bruza, P. D., Wang, Z., and Busemeyer, J. R. (2015). Quantum cognition: a new theoretical approach to psychology. Trends in Cognitive Sciences, 19(7):383–393.

9. Brysbaert, M., Mandera, P., and Keuleers, E. (2018). The word frequency effect in word processing: An updated review. Current Directions in Psychological Science, 27(1):45–50.

10. Buchsbaum, B. R. and D’Esposito, M. (2019). A sensorimotor view of verbal working memory. Cortex, 112:134–148.

11. Busemeyer, J. R. and Wang, Z. (2015). What is quantum cognition, and how is it applied to psychology? Current Directions in Psychological Science, 24(3):163–169.

12. Chen Zeng, T., Cheng, J. T., and Henrich, J. (2022). Dominance in humans. Philosophical Transactions of the Royal Society B, 377(1845):20200451.

13. Clark, A. (2013). Whatever next? predictive brains, situated agents, and the future of cognitive science. Behavioral and Brain Sciences, 36(3):181–204.

14. Clifton Jr, C., Ferreira, F., Henderson, J. M., Inhoff, A. W., Liversedge, S. P., Reichle, E. D., and Schotter, E. R. (2016). Eye movements in reading and information processing: Keith rayner’s 40 year legacy. Journal of Memory and Language, 86:1–19.

15. Conneau, A., Khandelwal, K., Goyal, N., Chaudhary, V., Wenzek, G., Guzmán, F., Grave, E., Ott, M., Zettlemoyer, L., and Stoyanov, V. (2019). Unsupervised cross-lingual representation learning at scale. arXiv preprint arXiv:1911.02116.

16. Crocker, M. W. (2012). Computational psycholinguistics: An interdisciplinary approach to the study of language. Springer Science & Business Media.

17. DeLong, K. A., Quante, L., and Kutas, M. (2014). Predictability, plausibility, and two late erp positivities during written sentence comprehension. Neuropsychologia, 61:150–162.

18. Demaree, H. A., Everhart, D. E., Youngstrom, E. A., and Harrison, D. W. (2005). Brain lateralization of emotional processing: historical roots and a future incorporating “dominance”. Behavioral and Cognitive Neuroscience Reviews, 4(1):3–20.

19. Demberg, V. and Keller, F. (2008). Data from eye-tracking corpora as evidence for theories of syntactic processing complexity. Cognition, 109(2):193–210.

20. Devlin, J., Chang, M.-W., Lee, K., and Toutanova, K. (2018). Bert: Pretraining of deep bidirectional transformers for language understanding. arXiv preprint arXiv:1810.04805.

21. Dirac, P. A. M. (1981). The principles of quantum mechanics. Number 27. Oxford University Press.

22. Engbert, R., Nuthmann, A., Richter, E. M., and Kliegl, R. (2005). Swift: a dynamical model of saccade generation during reading. Psychological Review, 112(4):777.

23. Frank, S. L. and Willems, R. M. (2017). Word predictability and semantic similarity show distinct patterns of brain activity during language comprehension. Language, Cognition and Neuroscience, 32(9):1192–1203.

24. Futrell, R., Gibson, E., and Levy, R. P. (2020). Lossy-context surprisal: An information-theoretic model of memory effects in sentence processing. Cognitive Science, 44(3):e12814.

25. Gibson, E. (1998). Linguistic complexity: Locality of syntactic dependencies. Cognition, 68(1):1–76.

26. Grave, E., Bojanowski, P., Gupta, P., Joulin, A., and Mikolov, T. (2018). Learning word vectors for 157 languages. In Proceedings of the International Conference on Language Resources and Evaluation (LREC 2018).

27. Gudder, S. P. (1970). A superposition principle in physics. Journal of Mathematical Physics, 11(3):1037–1040.

28. Guest, O. and Martin, A. E. (2021). How computational modeling can force theory building in psychological science. Perspectives on Psychological Science, 16(4):789–802.

29. Hahn, M., Futrell, R., Levy, R., and Gibson, E. (2022). A resource-rational model of human processing of recursive linguistic structure. Proceedings of the National Academy of Sciences, 119(43):e2122602119.

30. Hale, J. (2001). A probabilistic earley parser as a psycholinguistic model. In The Second Meeting of the North American Chapter of the Association for Computational Linguistics, Pittsburgh, Pennsylvania.

31. Hale, J. (2016). Information-theoretical complexity metrics. Language and Linguistics Compass, 10(9):397–412.

32. Hale, J., Lutz, D., Luh, W.-M., and Brennan, J. (2015). Modeling fmri time courses with linguistic structure at various grain sizes. In Proceedings of the 6th workshop on cognitive modeling and computational linguistics, pages 89–97.

33. Hale, J. T., Campanelli, L., Li, J., Bhattasali, S., Pallier, C., and Brennan, J. R. (2022). Neurocomputational models of language processing. Annual Review of Linguistics, 8:427–446.

34. Hohenstein, S., Laubrock, J., and Kliegl, R. (2010). Semantic preview benefit in eye movements during reading: A parafoveal fast-priming study. *Journal of Experimental Psychology: Learning*, Memory, and Cognition, 36(5):1150.

35. Hollis, G. and Westbury, C. (2016). The principals of meaning: Extracting semantic dimensions from co-occurrence models of semantics. Psychonomic Bulletin & Review, 23(6):1744–1756.

36. Huck, A., Thompson, R. L., Cruice, M., and Marshall, J. (2017). Effects of word frequency and contextual predictability on sentence reading in aphasia: An eye movement analysis. Aphasiology, 31(11):1307–1332.

37. Huettig, F. (2015). Four central questions about prediction in language processing. Brain Research, 1626:118–135.

38. Jensen, O., Pan, Y., Frisson, S., and Wang, L. (2021). An oscillatory pipelining mechanism supporting previewing during visual exploration and reading. Trends in Cognitive Sciences, 25(12):1033–1044.

39. Kennedy, A. and Pynte, J. (2005). Parafoveal-on-foveal effects in normal reading. Vision Research, 45(2):153–168.

40. Kennedy, A., Pynte, J., Murray, W. S., and Paul, S.-A. (2013). Frequency and predictability effects in the dundee corpus: An eye movement analysis. Quarterly Journal of Experimental Psychology, 66(3):601–618.

41. Kliegl, R., Grabner, E., Rolfs, M., and Engbert, R. (2004). Length, frequency, and predictability effects of words on eye movements in reading. European Journal of Cognitive Pyschology, 16(1/2):262–284.

42. Kliegl, R., Nuthmann, A., and Engbert, R. (2006). Tracking the mind during reading: the influence of past, present, and future words on fixation durations. Journal of Experimental Psychology: General, 135(1):12.

43. Koopmann, G. H., Song, L., and Fahnline, J. B. (1989). A method for computing acoustic fields based on the principle of wave superposition. The Journal of the Acoustical Society of America, 86(6):2433–2438.

44. Kuperberg, G. R. and Jaeger, T. F. (2016). What do we mean by prediction in language comprehension? Language, Cognition and Neuroscience, 31(1):32–59.

45. Kuperman, V., Schroeder, S., and Gnetov, D. (2023). Word length and frequency effects on text reading are highly similar in 12 alphabetic languages. https://psyarxiv.com/cbvjr/.

46. Kwon, N. and Sturt, P. (2014). The use of control information in dependency formation: An eye-tracking study. Journal of Memory and Language, 73:59–80.

47. Lee, M. D., Criss, A. H., Devezer, B., Donkin, C., Etz, A., Leite, F. P., Matzke, D., Rouder, J. N., Trueblood, J. S., White, C. N., et al. (2019). Robust modeling in cognitive science. Computational Brain & Behavior, 2:141–153.

48. Levy, R. (2008). Expectation-based syntactic comprehension. Cognition, 106(3):1126–1177.

49. Lewis, R. L., Vasishth, S., and Van Dyke, J. A. (2006). Computational principles of working memory in sentence comprehension. Trends in cognitive sciences, 10(10):447–454.

50. Li, X., Huang, L., Yao, P., and Hyönä, J. (2022). Universal and specific reading mechanisms across different writing systems. Nature Reviews Psychology, 1(3):133–144.

51. Lison, P., Tiedemann, J., and Kouylekov, M. (2018). Opensubtitles2018: Statistical rescoring of sentence alignments in large, noisy parallel corpora. In Proceedings of the 11th International Conference on Language Resources and Evaluation (LREC 2018). European Language Resources Association (ELRA).

52. Liu, H. (2008). Dependency distance as a metric of language comprehension difficulty. Journal of Cognitive Science, 9(2):159–191.

53. Loftus, G. R. (1985). Evaluating forgetting curves. *Journal of Experimental Psychology: Learning*, Memory, and Cognition, 11(2):397.

54. López-Peréz, P., Dampuré, J., Hernández-Cabrera, J., and Barber, H. (2016). Semantic parafoveal-on-foveal effects and preview benefits in reading: Evidence from fixation related potentials. Brain and Language, 162:29–34.

55. Malik-Moraleda, S., Ayyash, D., Gallée, J., Affourtit, J., Hoffmann, M., Mineroff, Z., Jouravlev, O., and Fedorenko, E. (2022). An investigation across 45 languages and 12 language families reveals a universal language network. Nature Neuroscience, 25(8):1014–1019.

56. Mascaro, O., Goupil, N., Pantecouteau, H., Depierreux, A., Van der Henst, J.-B., and Claidière, N. (2023). Human and animal dominance hierarchies show a pyramidal structure guiding adult and infant social inferences. Nature Human Behaviour, pages 1–13.

57. Mikolov, T., Grave, E., Bojanowski, P., Puhrsch, C., and Joulin, A. (2017). Advances in pre-training distributed word representations. arXiv preprint arXiv:1712.09405.

58. Mitchell, J. and Lapata, M. (2010). Composition in distributional models of semantics. Cognitive Science, 34(8):1388–1429.

59. Murre, J. M. and Dros, J. (2015). Replication and analysis of ebbinghaus’ forgetting curve. PloS One, 10(7):e0120644.

60. Niefind, F. and Dimigen, O. (2016). Dissociating parafoveal preview benefit and parafovea-on-fovea effects during reading: A combined eye tracking and eeg study. Psychophysiology, 53(12):1784–1798.

61. Norris, D. (2017). Short-term memory and long-term memory are still different. Psychological Bulletin, 143(9):992.

62. Oh, B.-D. and Schuler, W. (2023). Why does surprisal from larger transformer-based language models provide a poorer fit to human reading times? Transactions of the Association for Computational Linguistics, 11:336–350.

63. Pan, Y., Frisson, S., and Jensen, O. (2021). Neural evidence for lexical parafoveal processing. Nature Communications, 12(1):5234.

64. Pollatsek, A., Reichle, E. D., and Rayner, K. (2006). Tests of the ez reader model: Exploring the interface between cognition and eye-movement control. Cognitive Psychology, 52(1):1–56.

65. Postle, B. R. (2006). Working memory as an emergent property of the mind and brain. Neuroscience, 139(1):23–38.

66. Pothos, E. M. and Busemeyer, J. R. (2022). Quantum cognition. Annual Review of Psychology, 73:749–778.

67. Rayner, K., Warren, T., Juhasz, B. J., and Liversedge, S. P. (2004). The effect of plausibility on eye movements in reading. *Journal of Experimental Psychology: Learning*, Memory, and Cognition, 30(6):1290.

68. Reichle, E. D., Rayner, K., and Pollatsek, A. (2003). The ez reader model of eye-movement control in reading: Comparisons to other models. Behavioral and Brain Sciences, 26(4):445–476.

69. Reilly, R. G. and Radach, R. (2006). Some empirical tests of an interactive activation model of eye movement control in reading. Cognitive Systems Research, 7(1):34–55.

70. Roland, D., Yun, H., Koenig, J.-P., and Mauner, G. (2012). Semantic similarity, predictability, and models of sentence processing. Cognition, 122(3):267–279.

71. Rudin, W. (2017). Fourier analysis on groups. Courier Dover Publications.

72. Russo, A. G., De Martino, M., Mancuso, A., Iaconetta, G., Manara, R., Elia, A., Laudanna, A., Di Salle, F., and Esposito, F. (2020). Semantics-weighted lexical surprisal modeling of naturalistic functional mri timeseries during spoken narrative listening. Neuroimage, 222:117281.

73. Ryskin, R. and Nieuwland, M. S. (2023). Prediction during language comprehension: what is next? Trends in Cognitive Sciences.

74. Sayeed, A., Fischer, S., and Demberg, V. (2015). Vector-space calculation of semantic surprisal for predicting word pronunciation duration. In Proceedings of the 53rd Annual Meeting of the Association for Computational Linguistics and the 7th International Joint Conference on Natural Language Processing (Volume 1: Long Papers), pages 763–773.

75. Schotter, E. R., Angele, B., and Rayner, K. (2012). Parafoveal processing in reading. *Attention, Perception*, & Psychophysics, 74(1):5–35.

76. Schrimpf, M., Blank, I. A., Tuckute, G., Kauf, C., Hosseini, E. A., Kanwisher, N., Tenenbaum, J. B., and Fedorenko, E. (2021). The neural architecture of language: Integrative modeling converges on predictive processing. Proceedings of the National Academy of Sciences, 118(45):e2105646118.

77. Schwering, S. C. and MacDonald, M. C. (2020). Verbal working memory as emergent from language comprehension and production. Frontiers in Human Neuroscience, 14:68.

78. Shliazhko, O., Fenogenova, A., Tikhonova, M., Mikhailov, V., Kozlova, A., and Shavrina, T. (2022). mgpt: Few-shot learners go multilingual. arXiv preprint arXiv:2204.07580.

79. Siegelman, N., Schroeder, S., Acartürk, C., Ahn, H.-D., Alexeeva, S., Amenta, S., Bertram, R., Bonandrini, R., Brysbaert, M., Chernova, D., et al. (2022). Expanding horizons of cross-linguistic research on reading: The multilingual eye-movement corpus (meco). Behavior Research Methods, pages 1–21.

80. Snell, J. and Grainger, J. (2019). Readers are parallel processors. Trends in Cognitive Sciences, 23(7):537–546.

81. Staub, A. (2015). The effect of lexical predictability on eye movements in reading: Critical review and theoretical interpretation. Language and Linguistics Compass, 9(8):311–327.

82. Sun, K. (2023). Attention-aware semantic relevance for predicting chinese sentence reading. Manuscript.

83. Sun, K., Wang, Q., and Lu, X. (2023a). An interpretable measure of semantic similarity for predicting eye movements in reading. Psychonomic Bulletin & Review, pages 1–16.

84. Sun, K., Wang, R., and Baayen, H. (2023b). Attention-aware measures of semantic relevance for predicting human reading behavior. Linguistics.

85. Ueno, T., Saito, S., Rogers, T. T., and Ralph, M. A. L. (2011). Lichtheim 2: synthesizing aphasia and the neural basis of language in a neurocomputational model of the dual dorsal-ventral language pathways. Neuron, 72(2):385–396.

86. Vasilev, M. R. and Angele, B. (2017). Parafoveal preview effects from word n+ 1 and word n+ 2 during reading: A critical review and bayesian metaanalysis. Psychonomic Bulletin & Review, 24:666–689.

87. Vaswani, A., Shazeer, N., Parmar, N., Uszkoreit, J., Jones, L., Gomez, A. N., Kaiser, Ł., and Polosukhin, I. (2017). Attention is all you need. Advances in Neural Information Processing Systems, 30.

88. Veldre, A. and Andrews, S. (2016). Is semantic preview benefit due to relatedness or plausibility? Journal of Experimental Psychology: Human Perception and Performance, 42(7):939.

89. Wen, Y., Snell, J., and Grainger, J. (2019). Parallel, cascaded, interactive processing of words during sentence reading. Cognition, 189:221–226.

90. Wilcox, E. G., Gauthier, J., Hu, J., Qian, P., and Levy, R. (2020). On the predictive power of neural language models for human real-time comprehension behavior. arXiv preprint arXiv:2006.01912.

91. Wood, S. N. (2017). Generalized Additive Models: An Introduction with R. Chapman and Hall/CRC.

